# The hematopoietic landscape at single-cell resolution reveals unexpected stem cell features in naked mole-rats

**DOI:** 10.1101/859454

**Authors:** Stephan Emmrich, Marco Mariotti, Masaki Takasugi, Maggie E. Straight, Alexandre Trapp, Vadim N. Gladyshev, Andrei Seluanov, Vera Gorbunova

## Abstract

Naked mole-rats are the longest-lived rodents endowed with resistance to cancer and age-related diseases, yet their stem cell characteristics remain enigmatic. We profiled the naked mole-rat hematopoietic system down to single-cell resolution, and identified several unique features likely contributing to longevity. In adult naked mole-rats red blood cells are formed in spleen and marrow, a neotenic feature beneficial for hypoxic environments and to prevent anemia. Platelet numbers are lower compared to short-lived mice, which may preclude age-related platelet increase and thrombosis. T cells mature in thymus and lymph nodes, providing a supply of T cells after age-related thymus involution. The pool of quiescent stem cells is higher than in mice, and HSCs overexpress an oxidative phosphorylation signature, revealing a new paradigm of stem cell metabolism to benefit longevity and oppose oncogenesis. Our work provides a platform to study immunology and stem cell biology in an animal model of healthy aging.

**HIGHLIGHTS:**

- Flow cytometry labelling panel to purify viable naked mole-rat HSPCs
- The spleen as the major site of erythropoiesis in the naked mole-rat
- Naked mole-rats show extrathymic T-cell development under homeostatic conditions
- Naked mole-rat hematopoietic stem cells (HSCs) have high OXPHOS activity

## INTRODUCTION

The naked mole-rat (*Heterocephalus glaber*) has become an attractive animal model in aging research due to its longevity and resistance to disease (Gorbunova et al., 2014). This remarkable rodent with the size of a mouse has a maximum lifespan of over 30 years in captivity and does not display increased mortality during aging (Ruby et al., 2018). Naked mole-rat cells feature higher translational fidelity due to split processing of 28S rRNA (Azpurua et al., 2013), express a unique splicing product from the senescence-inducing INK4/ARF locus and produce abundant high molecular weight hyaluronic acid responsible for the resistance of naked mole-rats to solid tumors (Tian et al., 2013; Tian et al., 2015). Physiological traits include an extreme resistance to anoxia through fructose metabolism (Park et al., 2017), skin acid insensitivity and a serum metabolome reflecting calorically restricted mice (Lewis et al., 2018; Smith et al., 2011).

The blood system is the most regenerative tissue, producing >10^14^ cells per year in humans (Dancey et al., 1976). Fostered by recent advances in single-cell transcriptomics (Paul et al., 2015), hematopoiesis is viewed as a continuum of individual cells that traverse the differentiation process from unprimed hematopoietic stem and progenitor cells (HSPCs) directly into unipotent progenitors (Velten et al., 2017). Studies of the unperturbed hematopoietic system are most advanced in their understanding of HSPC hierarchies and concepts of stemness in mouse (Laurenti and Gottgens, 2018). There are, however, fundamental differences in certain aspects of the blood system between mice and humans, reviewed in (Copley and Eaves, 2013). At the genetic level, orthologs for one of the major murine HSC markers, Sca-1 (Ly6a), are found only in rodents but not in primates, sauropsida, carnivores or fish. Interestingly, naked mole-rats are among the few rodents without a Sca-1 ortholog. Considering its for longevity we set out to characterize the hematopoietic system of the naked mole rat and develop a platform for future studies in this long-lived rodent.

We developed a flow cytometry (FACS) labelling strategy using cross-reactive antibodies to sort, culture and transplant naked mole-rat hematopoietic stem and progenitor cells (HSPCs). A panel of six surface markers purified distinct HSPC fractions. Single-cell transcriptomics revealed conservation of a multipotent progenitor (MPP) stage primed towards lymphomyeloid commitment and a megakaryocytic-erythroid progenitor (MEP) stage marked by coordinated GATA1/2 expression. Comparative analysis of stem and progenitor hierarchies of 3 species delineated high OXPHOS signaling and several upregulated cytoprotective mechanisms in naked mole-rat HSPCs. We found further adaptations to longevity in the hematopoietic system of naked mole-rats such as a substantially altered T cell (TC) pool, extrathymic TC maturation and splenic erythropoiesis, revealing systemic deviations from traditional concepts of hematopoiesis some of which have neotenic features and may together promote longevity. Our findings provide a comprehensive resource for the studies of immunology, inflammation and stem cell biology in the naked mole-rat as a model of healthy aging.

## RESULTS

### Cross-reactive CD antibodies for separation of naked mole-rat hematopoietic cells

Despite recent advances in understanding the mechanisms longevity and disease resistance in naked mole-rats, little is known regarding stem cells and the hematopoietic system in these animals. We screened commercially available monoclonal antibodies (MoAb) to CD markers in human, mouse, rat and guinea pig. We identified human CD11b, CD18, CD34 and CD90, mouse CD11b and CD125, rat Thy1.1 and guinea pig CD45 cross-reactive to bind distinct subsets of viable naked mole-rat bone marrow (BM) cells (Figure 1A). Specific binding was validated by light scatter properties of the marker-positive populations (Figure S1A-E). We then used these antibodies to further characterize hematopoietic cell subsets in the naked mole-rat.

**Figure 1.**
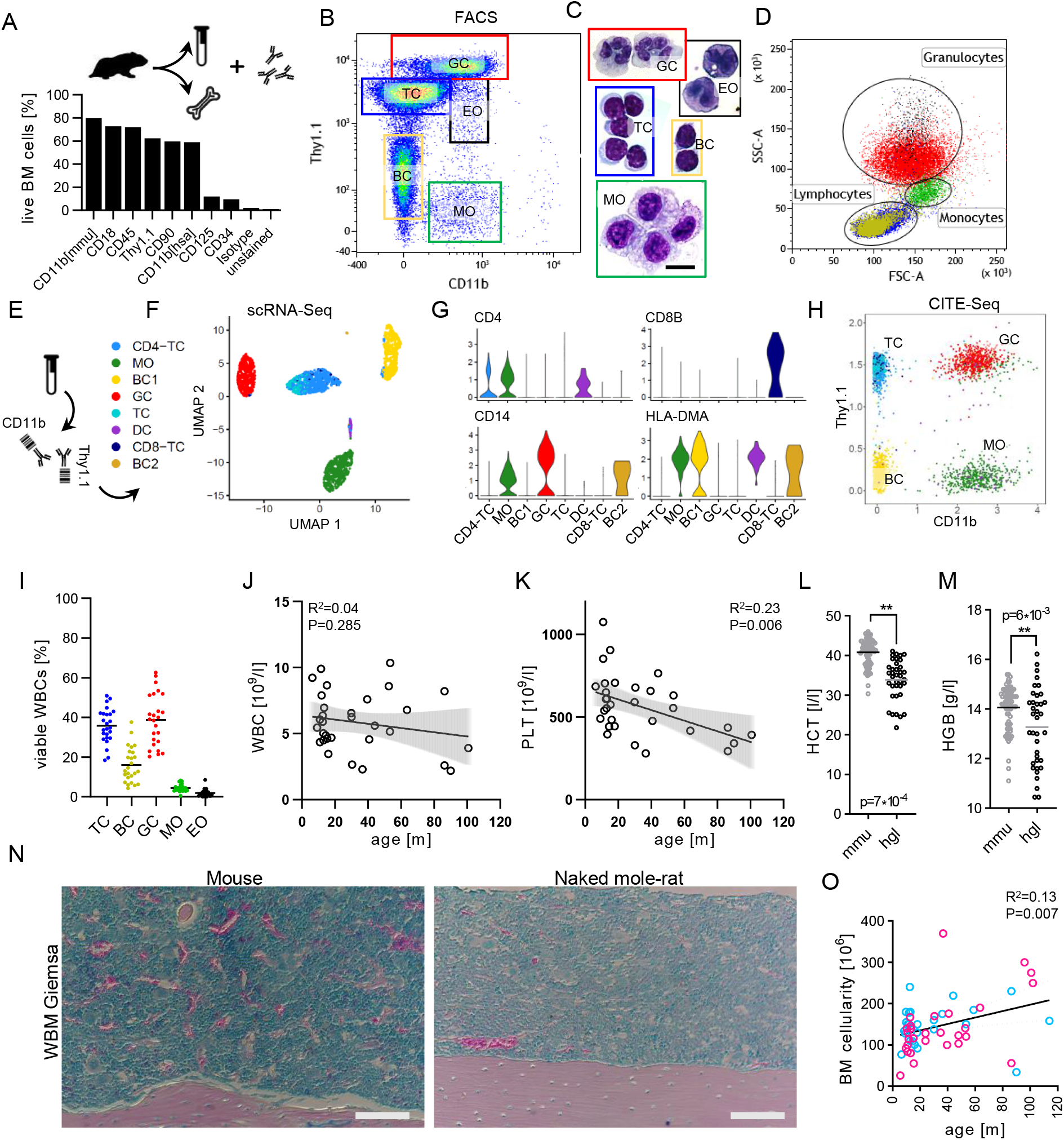
The blood cell composition of naked mole-rats is different from mice. (A) Frequency of naked mole-rat BM cells stained with cross-reactive antibodies. (B) Representative FACS gating of blood stained with Thy1.1 and CD11b. (C) May-Grünwald-Giemsa staining of cells sorted from respective blood FACS gates; Scale bar 20µm, same magnification for each micrograph. (D) FACS backgating of each fraction from (B) into side (SSC) and forward (FSC) scatters. Major blood cell type gates for lymphocytes, monocytes and granulocytes were gated on viable singlets as shown in Figure S1C. (E) Experimental design of blood CITE-Seq using antibodies from (B). (F) UMAP of Louvain-clustered blood single-cell transcriptomes, color legend is used throughout scRNA-Seq data in Figures 1, S1. (G) Expression of selected genes across blood single-cell clusters. Expression level, scaled UMI counts per cell. (H) CITE-Seq scaled UMI counts per cell for Thy1.1 and CD11b, colored with reference to (B). (I) Frequencies of normal naked mole-rat blood cell fractions gated as in (B); WBCs, white blood cells; N=25. Bloodcounter volumetric leukocytes (J) and platelet (K) numbers across animal age. R^2^, Pearson correlation coefficient; p-values were determined by conventional linear regression fitting both slope and intercept; N=31. Bloodcounter hematocrit percentage (L) and hemoglobin concentration (M) between normal mouse [mmu] and naked mole-rat [hgl] blood. p-values were determined by unpaired Welch’s t-test; N(mmu)=81, N(hgl)=37. (N) Giemsa staining of femur sections, medullary canal of diaphysis; Scale bar 100µm. (O) BM cellularity as determined by Trypan-Blue counting of RBC-lysed BM over animal age; N=54; pink, female [25]; blue, male [29]; R^2^, Pearson correlation coefficient; p-value determined by linear regression. See also Table S1, Figure S1-2.

### The naked mole-rat hemogram differs from human and mouse

When we stained red blood cell (RBC) depleted naked mole-rat peripheral blood with Thy1.1 and CD11b, the 5 major blood cell types could be distinguished: neutrophil granulocytes (GC), eosinophil granulocytes (EO), T cells (TC), B cells (BC) and monocytes (MO) (Figure 1B). Cytochemistry of FACS-purified cells revealed multi-lobulated nuclei and a pH-neutral cytoplasm for Thy1.1^+++^/CD11b^+^ GCs and bi-lobulated nuclei with a high density of acidic granulation for Thy1.1^++^/CD11b^+^ EOs, both populations exhibiting granulocytic scatter properties (Figure 1C-D). Two populations with lymphocytic morphology and size are labelled as Thy1.1^++^/CD11b^−^ and Thy1.1^+^/CD11b^−^, while Thy1.1^−^/CD11b^+^ cells resemble MOs (Figure 1B-D).

To increase cell type resolution we performed CITE-Seq (Stoeckius et al., 2017) for CD11b and Thy1.1 on sorted naked mole-rat blood cell populations (Figure 1E, S1F). A *de novo* transcriptome assembly from deep sampling of naked mole-rat whole marrow was prepared according to the FRAMA pipeline (Bens et al., 2016) and used for transcript annotation, which revealed hundreds of previously unannotated genes as well as thousands of novel transcript isoforms (Table S1, Figure S1G-J). We clustered 2594 cells into 8 distinct cell types (Figure 1E-F). A common blood TC signature comprising CD3 homologues, LAT, LCK and CD27 was expressed in CD4-TCs, CD8-TCs and a TC subset lacking a functional identification marker (Figure S1K-L). While CD8-TCs were rare (1.4%), and CD4-TCs constituted the majority of T cells (23.8%), only a subset of CD4-TCs expressed CD4 mRNA (Figure 1G). In fact, MO and dendritic cell (DC) clusters had higher fractions of CD4-expressing cells. The cryptic TC cluster (6.9%) lacked expression of genes detected in both CD4- and CD8-TCs, such as CCR7 (Figure S1L), and did not express canonical markers for regulatory T cells (no orthologue for CD25, FOXP3 not expressed) or Natural Killer cells (CD56, KLRK1, LY94 not expressed, no orthologue for NKp44). However, while NCR3 was expressed by all naked mole-rat blood cell types, CD8-TCs specifically expressed NKG7, Granzyme A and TARP (Wolfgang et al., 2000), an alternate reading frame of the TC receptor γ locus produced in non-lymphoid tissues and prostate or breast cancers (Figure S1L-M).

While both TCs and BCs were not labelled by CITE-CD11b, all three myeloid clusters were CITE-CD11b^+^ (Figure 1H). DCs expressed CD4, CD86, BATF3 and several HLA-D orthologues (Figure 1G, S1H). FACS-based quantitation of naked mole-rat blood cells was corroborated by blood counter measurements (Figure S1M).

It is well described that murine blood leukocyte counts increase with age (Figure S2A). This was not observed in a naked mole-rat cohort spanning less than 1 to 8 years of age (Figure 1J). Likewise, blood platelet levels increased in mice, but did not increase and were ∼2-fold lower in naked mole-rats (Figure 1K, S2B-D). The hematocrit was markedly lower for naked mole-rats (33.8%) than mice (40.8%), moreover RBC hemoglobin was slightly decreased in naked mole-rats (Figure 1L-M). Total RBC numbers did not change with age (Figure S2E-F).

Thus, the unique features of naked mole-rat blood are a definitive TC pool wherein most cells are CD4^−^/CD8^−^, pointing towards altered immune functions as an adaptation to longevity. Moreover, naked mole-rats did not display age-associated increase of blood leukocytes and platelets. This may reduce chronic inflammations and prevent age-associated thrombosis.

### Characterization of the stem cell compartment in the BM

The mammalian marrow is the primary site of hematopoiesis, harboring HSCs and producing RBCs and leukocytes (Eaves, 2015). Imaging of longitudinal femur sections showed fewer erythropoietic islets and Megakaryocytes (MKs) for naked mole-rat long bones as compared to mice (Figure 1N, S2G). While BM cellularity in mice increases during very early life stages and starts to decline at the onset of middle age (Colvin et al., 2004), we observed a slight increase of BM cellularity over the first 10 years of age in naked mole-rats (figure 1O, S7C).

One of the first milestones towards prospective isolation of HSCs was the early notion that the cell fraction with hematopoietic regenerative potential was nearly devoid of markers of mature blood cell types (Muller-Sieburg et al., 1986). We thus designed a naked mole-rat lineage depletion cocktail (LIN) from the cross-reactive antibodies which we identified (Figure 1A). CD11b and CD18 both form the integrin Mac-1, marking myeloid or NKC commitment. CD125, the IL-5 receptor alpha subunit, is primarily expressed on eosinophils and activated BCs (Huston et al., 1996). CD34 and CD90/THY1 are human stem cell markers (Bhatia et al., 1997). However, we found that the Thy1.1 MoAb labels additional cells not commonly stained with CD90 MoAb in naked mole-rat BM, most likely due to different epitopes, each with a proteoform-specific label for naked mole-rat THY1 (Figure S2H). Scatter backgating demonstrated that CD90^++^ cells corresponding to Thy1.1^+++^ cells were neutrophils, while dim CD90^+^ cells were TCs. We therefore used CD90 antigen to deplete committed cells from the Thy1.1 label, completing our LIN to CD11b/CD18/CD90/CD125 (Figure 2A). Given that this LIN lacks definitive depletion markers for pan BCs, NKCs and erythroid precursors we set the LIN^−^ gate at ∼15% of viable WBCs, which purified HSPCs as demonstrated by significantly higher colony formation than LIN^+^ or BM (Figure 2B). The LIN^+^ fraction enriched GC, TC and BC populations (Figure 2C). Most cells of the LIN^−^ fraction were Thy1.1^−^/CD34^−^ (CP7; candidate population), resembling committed cells not covered by the naked mole-rat LIN cocktail (Figure 2D). Surprisingly, both LIN^+^ and LIN^−^ contain a Thy1.1^++^/CD34^++^ population, which we termed CP1 and CP2, respectively. CP3 is LIN^−^/Thy1.1^+^/CD34^++^, CP4-5 are Thy1.1^−^ with decreasing CD34, while CP6 resembles blood BCs (Figure 2D). Checking which LIN factor is differentially expressed on Thy1.1^++^/CD34^++^ cells we found CD11b and CD90 absent in CP7 and CP2 but present in CP1 and in most viable Thy1.1^++^/CD34^++^ cells, all four being negative for CD125 (Figure 2F-H). We subset distinct cell populations of the LIN^−^ fraction to find that only CP1, CP2 and CP3 are <1% of total BM leukocytes (Figure 2E), the common frequency of a hematopoietic stem cell compartment (Morrison and Weissman, 1994).

**Figure 2.**
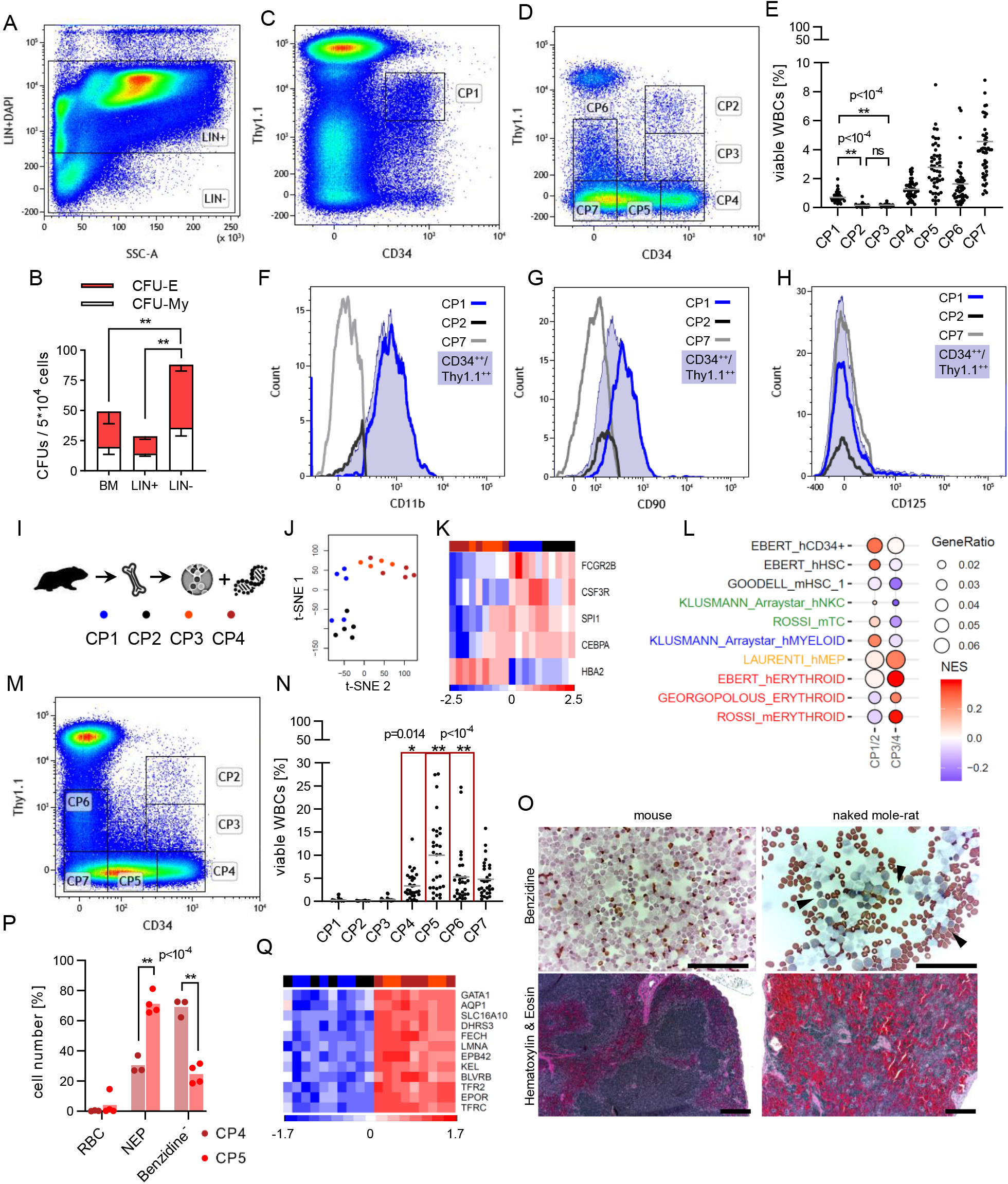
Normal erythropoiesis predominantly occurs in the spleen. Sorting strategy for the HSPC compartment with lineage (LIN) depletion (A), gating of LIN^+^ CP1 (C) and gating of LIN^−^ CP2-7 (D). (B) Colony assay grown at 37°C; CFU-E, colony forming unit erythroid; CFU-My, colony forming unit myeloid. Error bars denote s.d., p-value determined by Sidak’s Two-way ANOVA; N=4. (E) Frequencies of BM CP cell fractions. p-value determined by Brown-Forsythe’s One-way ANOVA; N=48. Signal intensities of CD11b (F), CD90 (G) and CD125 (H) of indicated BM cell fractions; CD34^++^/Thy1.1^++^ [steelblue AUC] not LIN gated. (I) Workflow for bulk RNA-Seq with color legend of analyzed cells. (J) Unsupervised t-SNE clustering; vst-transformed counts as input. (K) Clustering by Euclidean distance using indicated genes; voom-transformed and row-scaled expression values. (L) ssGSEA of combined CP1/2 and CP3/4 groups showing top 10 q-value genesets; each q < 10^-10^. NES, normalized enrichment score; GeneRatio, (signature ∩ term) / (signature ∩ all terms). (M) LIN^−^ fraction of the spleen with sorting gates analog to BM. (N) Frequencies of spleen CP cell fractions; N=30. p-value determined by Sidak’s Two-way ANOVA comparing BM vs spleen. (O) Benzidine (top) staining of whole spleen cytospins [scale bar 250µm; arrows indicate NEPs] and H&E (bottom) staining of spleen sections [scale bar 200µm]. (P) Relative counts of Benzidine-stained cytospins of sorted spleen fractions. p-value determined by Sidak’s Two-way ANOVA comparing BM vs spleen; N=4. NEP, nucleated erythroid progenitors. (Q) Cell clustering by Euclidean distance using top 12 MEP/erythroid leading edge genes. See also Figure S2.

We next performed population RNA-Sequencing of the four least abundant CD34^+^ BM fractions to annotate the developmental status (Figure 2I). We detected 9531 naked mole-rat orthologs expressed across 19 samples (Figure S2I-J, Table S2). Unsupervised clustering by t-distributed stochastic neighborhood embedding (t-SNE) separated CP1/CP2 from CP3/CP4, however, two CP4 samples cluster with the CP3 group and two CP1 samples cluster in the CP2 group (Figure 2J).

Our reanalysis of a human and a murine HSPC RNA-Seq dataset revealed that sharpest clustering differences were obtained between erythroid and myeloid lineages (Figure S2K-T). We thus probed the expression of naked mole-rat othologs HBA2 vs CD32 (FCGR2B) and the G-CSF receptor (CSF3R) and saw an inverse correlation between the globin gene and the two surface markers instructive for myeloid development (Figure 2K). Indeed, single sample gene set enrichment analysis (ssGSEA) associated MEP and erythroid gene expression modules to clusters CP3/4, while assigning stem cell, lymphoid and myeloid identity to clusters CP1/2 (Figure 2L).

Altogether, we have developed a MoAb-based labelling panel for the primitive HSPC compartment, wherein diminished Thy1.1 expression of CD34^+^ cells correlated with erythroid fate decision.

### The naked mole-rat spleen is the major site of erythropoiesis

In most mammals the spleen primarily acts as to recycle aged erythrocytes (Mebius and Kraal, 2005). However, species-specific adaptations have been found, such as the murine spleen acting as a reservoir of MOs or the equine spleen as a storage of up to 30% RBCs (Kunugiyama et al., 1997; Swirski et al., 2009). We saw a drastic difference in the LIN staining pattern as compared to BM with a strongly expanded LIN^dim^ population corresponding to Thy1.1^−^/CD34^−/+/++^ cells in naked mole-rat spleens (Figure S2U-V). The frequencies of CP4 (2.3-fold), CP5 (3.5-fold) and CP6 were significantly increased in naked mole-rat spleens relative to BM (Figure 2M-N). Since the CP4 immunophenotype Thy1.1^−^/CD34^++^ featured erythroid gene expression, we found increased RBC content in splenic organ sections compared to mice (Figure 2O). Moreover, Benzidine-stained cytospins of whole spleens revealed significantly more RBCs in naked mole-rats (Figure 2O, S2W). We further detected Benzidine^+^ nucleated erythroid precursors in naked mole-rat but not mouse spleens (Figure 2O). Strikingly, in adult mice where normal erythropoiesis is known to occur in the BM, the number of nucleated erythroid progenitors diminishes 19-fold from BM to Spleen, in contrast to a 2-fold increase from BM to Spleen in naked mole-rats (Figure S2W), pointing towards shared splenic and medullary erythropoiesis. To link elevated nucleated erythroid progenitor levels with expansion of the Thy1.1^−^/CD34^+/++^ compartment we sorted CP3 and CP4 cells from spleens for Benzidine staining (Figure 2P). This clearly demonstrated an increase of nucleated erythroid progenitors along with a decline of CD34 expression from CP4 to CP5. We thus defined erythroid cell fate in the naked mole-rat LIN^−^ compartment by a gradual loss of Thy1.1, directly followed by successive downregulation of CD34. Commitment onsets at CP4, marked by elevated expression of GATA1, EPOR, TFRs, KEL and FECH in CP3/4 (Figure 2Q).

In summary we have mapped sequential erythroid progenitor populations and revealed the spleen as a site of erythropoiesis. This represents a neotenic trait, as in other mammals splenic hematopoiesis takes place during fetal development. In the naked mole rat splenic hematopoiesis may compensate for lower BM cellularity and thick-walled bones (Figure S7D). Furthermore, shared splenic and medullary erythropoiesis may have evolved as an adaptation to hypoxia, however, it may also benefit longevity by preventing age-associated anemia.

### Functional evidence for LTCs (LIN^−^/Thy1.1^++^/CD34^++^) as naked mole-rat HSCs

To our surprise the two master regulator transcription factors of myeloid differentiation, PU.1 (SPI1) and CEBPα, are upregulated in the majority of CP3 samples along with CP1/2 (Figure 2K). Revisiting the primary clustering with a trimmed dataset enabled recapitulation of FACS groupings (Figure 3A, S3A, Table S2). Strikingly, the strongest association for CP3 was MEPs (Figure 3B). CP1 was enriched with the most HSC-associated genesets (Figure 3B). The human CD34+ signature displayed a NES gradient from CP1 to CP4, revealing a Mac-1^+^ HSPC fraction with a stemness expression profile in adult naked mole-rats. Moreover, CP1-specific genes are shared with lymphoid-primed multipotent progenitors (LMPPs) and the highest gene ratio is seen for CMP (common myeloid progenitor) gene expression (Figure 3B). By contrast the top scoring genesets for CP2 cells were downregulation of cell division and moderate expression of human thymic early TC progenitors (hETP; Figure 3B).

**Figure 3.**
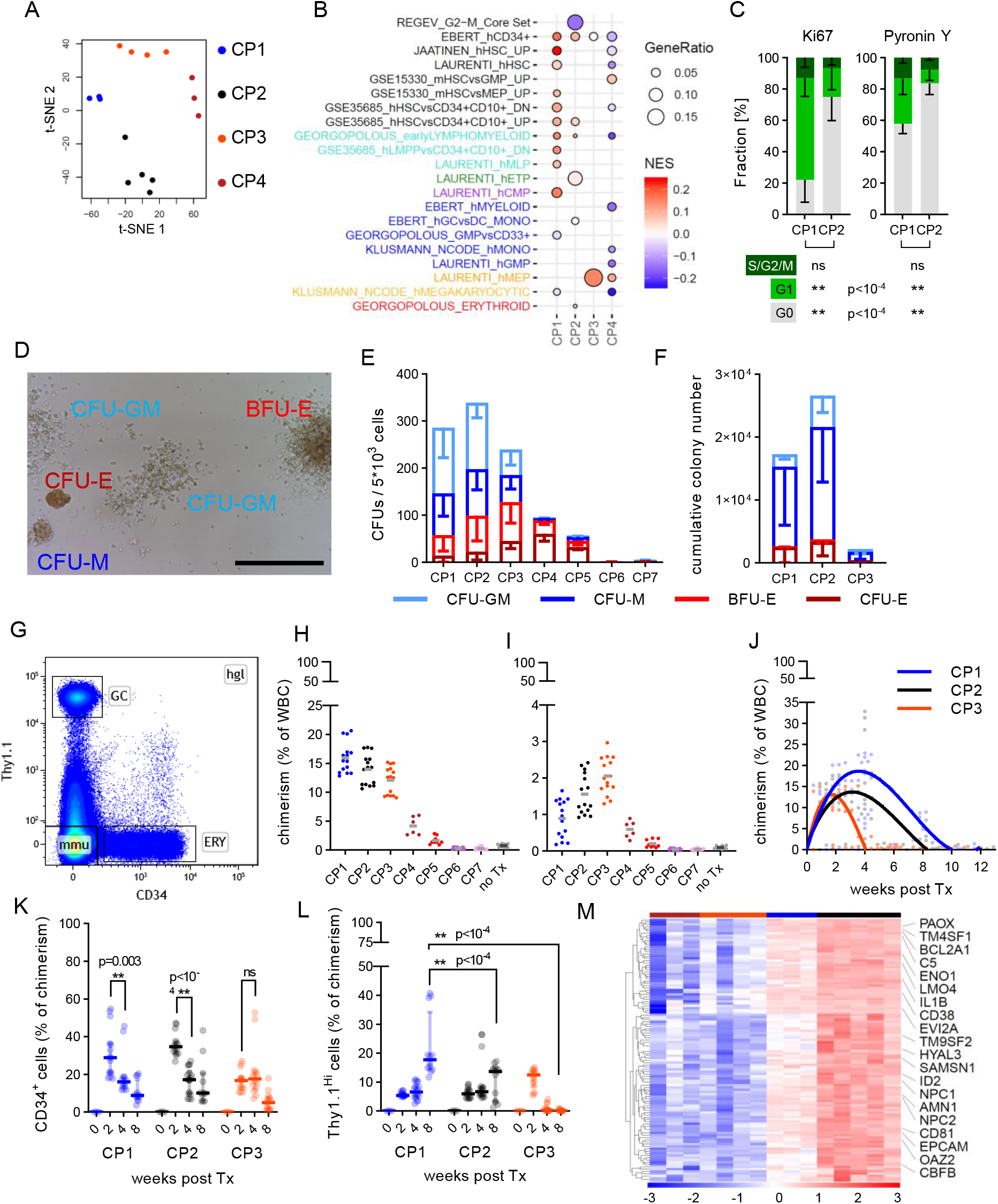
LTCs define the primitive HSC compartment. (A) Unsupervised t-SNE clustering using a trimmed datased from Figure 2J. (B) ssGSEA showing highest 30% GeneRatio genesets; each q < 0.05. (C) Cell cycle staining with Ki67 (left; N=12) or Pyronin Y (right; N=6); p-value determined by Sidak’s Two-way ANOVA. (D) Representative image of colony morphologies grown at 32°C. BFU-E, burst forming unit erythroid; CFU-M, colony forming unit macrophage; CFU-GM, colony-forming unit granulocyte-monocyte; Scale bar 250µm. Colony assay (E; N=5) or Replated assays (F; N=3). Error bars denote s.d., p-value determined by Sidak’s Two-way ANOVA. (G) Gating strategy to quantify total engraftment; hgl, naked mole-rat cells; mmu, mouse cells; ERY, CD34^+^ erythroid xenograft; GC, Thy1.1^+++^ granulocytic xenograft. NSGS chimerism 2 weeks post Tx in BM (H) or spleen (I); total recipients from 3 donors for each CP graft. (J) BM chimerism over time; recipients from 5 donors for weeks 4,8 and 12. p-value determined by Fisher’s Two-way ANOVA; curve-fitting by cubic polynomial. (K) ERY proportion of engraftment; p-value determined by Tukey’s Two-way ANOVA. (L) GC proportion of engraftment; p-value determined by Tukey’s Two-way ANOVA. (M) 116 gradually downregulated genes from CP2 to CP1, displayed are 20 genes with known roles in hematopoiesis. See also Figure S3-4.

Hence, we tested whether our ssGSEA modelling correctly segregated CP2 as non-cycling cells. A cross-reactive Ki67 MoAb revealed a typical staining pattern in naked mole-rat WBM (Figure S3B). Alternatively, we applied Pyronin Y staining to corroborate the cell cycle status in viable cells (Figure S3C). Both methods clearly demonstrated more cells in G_0_ for CP2 than CP1 (Ki67 3.4-fold, Pyronin Y 1.5-fold; Figure 3C).

The capacity to give rise to several distinct lineages via differentiation, referred to as multipotency, can be assayed through quantitation of progenitor frequencies during colony formation (Roy et al., 2012). We observed that naked mole-rat HSPCs grew best at 32°C in methylcellulose supplemented with human cytokines (Figure 3D). Scoring of colony-forming unit (CFU) types was validated by cytochemistry of picked colonies (Figure S3D-F). Out of all naked mole-rat BM populations only CP6 and CP7 did not grow in methylcellulose assays (Figure 3E). Furthermore, the proportion of erythroid over total colonies declined from CP4/5 (0.94/0.81) to CP3 (0.53), CP2 (0.29) and was lowest in CP1 (0.2) (Figure 3E). Myeloid output was not significantly different between CP1 and CP2 but decreased in CP3. Serial replating yielded 1.5-fold more total colonies for CP2 compared to CP1, although no colony type frequency was significantly altered between these two, as seen for original platings (Figure 3F). Consistent with ssGSEA annotation, Benzidine-stained colony assays showed an increase in the proportion of hemoglobin-containing colonies from CP1 (0.08) over CP2 (0.26) and CP3 (0.56) to CP4/5 (0.94/0.79; Figure S3G). Notably CP1 colonies featured less mixed Benzidine^+/–^ colonies than CP2, pointing towards lymphomyeloid lineage restriction of CP1 cells.

Another hallmark of adult stem cells is self-renewal to perpetuate the stem cell pool under homeostatic conditions. The gold standard assay for HSC self-renewal are serial transplantations, and although human xenotransplantation grafts generally perform poor in secondary recipients, high levels of sustained primary engraftments could be obtained in a variety of immunodeficient mouse models (Goyama et al., 2015). Given the successful *in vitro* growth of naked mole-rat HSPCs with human cytokines we reasoned that the NSGS mouse model with constitutive production of human IL-3, M-CSF and SCF would render optimal support to naked mole-rat xenografts (Wunderlich et al., 2010). We indeed observed robust engraftment rates for CP2 at 4 weeks post transplantation in recipient BM as compared to untransplanted mice (Figure 3G, S3H). Xenografts recapitulated the Thy1.1/CD34 FACS staining pattern of naked mole-rat BM origin and were quantifiable (Figure S3I-L). At week 2 host BM chimerism resembled colony yields from methylcellulose assays with CP6/7 engraftments below background, supporting the notion that the naked mole-rat HSPC compartment is CD34^+^ as in humans (Figure 3H). All other populations produced clearly detectable naked mole-rat engraftment in NSGS mice ranging from 1.6% (CP5) over 4.2% (CP4) and 12.1% (CP3) to 14% (CP2) and 16% (CP1). Repopulation of host spleens was markedly reduced for all engrafted groups; Strikingly CP3 was functionally classified as the most primitive committed erythroid progenitor and enriched in naked mole-rat spleens with higher engraftment than CP2 (Figure 3I). Though FACS verified xenograft cells in blood for CP1, CP2 and CP3, levels ranged below 1% of viable leukocytes (data not shown). BM engraftment at week 4 for CP3 (1%) depleted earlier than for CP1 (12.8%, p<10^-4^) and CP2 (11.7%, p<10^-4^) (Figure 3J). The early loss of erythroid-primed CP3 is consistent with higher residual chimerism at week 8 for myeloid-primed CP1 compared to CP2 (5.3% vs 1.3%, p=0.04). Unexpectedly none of the three most primitive stem and progenitors sustained BM engraftment past 12 weeks, a fact we primarily attribute to the difference in body temperature between naked mole-rats (thermoneutral at ∼32°C) and humans or mice (Figure 3J). Next we quantified lineage commitment over time by selecting xenograft CD34^+^ cells as measure for erythropoiesis. Interestingly, while CP1 and CP2 CD34^+^ cells decline towards week 4, CP3 CD34^+^ output remained similar (Figure 3K). Conversely, we used Thy1.1^+++^ cells as myeloid output, which revealed most efficient myelopoiesis at week 8 in CP1 compared to CP2 and CP3 (Figure 3L). As a surrogate for self-renewal we quantified the complete Thy1.1^+^/CD34^+^ compartment of xenografts from which all three populations originated (Figure S3M). Although this rapidly depleted for all cell types at week 4, the initial replicative burst was greater in CP1 compared to CP2 (Figure S3N). This is consistent with our cell cycle measurements wherein CP2 showed a higher fraction of resting cells (Figure 3C). Finally, we co-stained transplanted BM from week 2 with hCD11b, a MoAb reactive with naked mole-rat but not binding to mouse cells, and CD45.1, which labels all hematopoietic cells in NSGS mice, and gated a selection hCD11b^−^ and hCD11b+ (Figure S4A). This clearly showed that CP2 maintained its original <u>L</u>IN^−^/<u>T</u>hy1.1^++^/<u>C</u>D34^++^ LTC population and gave rise to CD11b^+^/Thy1.1^++^/CD34^++^ cells, whereas CP1 expanded CD11b^+^/Thy1.1^++^/CD34^++^ yet failed to form a CP2 population *in vivo* (Figure S4A-D).

Altogether CP1 cells strongly resemble MPPs expressing Mac-1, CD90 and a stemness signature. Although we cannot exclude the possibility that CP1 contains activated HSCs, all evidence points towards a primitive multipotent leukocyte progenitor with severely decreased capacity of differentiating towards the erythroid lineage. Moreover, we functionally defined the most primitive HSC compartment as LTCs, exhibiting the highest degree of quiescence, multipotency and self-renewal.

### LTCs and CP1 MPPs share a homogenous cell population with HSC expression profile

HSCs are known to be at the apex of the hematopoietic hierarchy, thus we checked which genes were successively downregulated during transition from LTCs to CP1 and CP3/4 (Figure 3M). We retrieved 116 genes showing such expression pattern, of which 40 are found in the specific LTC signature (Figure S3A, Table S2). A key finding was high expression of ID2, which blocks BC differentiation, can enhance erythropoiesis and expands HSCs (Ji et al., 2008; van Galen et al., 2014). This population also showed high expression of CD81, a tetraspanin which has been shown to maintain self-renewal in HSCs (Lin et al., 2011).

To further increase the resolution of the naked mole-rat HSPC compartment we sorted and pooled BM CP1-7 cells for CITE-Seq (Figure S4D-E). 11669 orthologs and the 3 CITE-signals were mapped to 12914 cells, yielding 16 clusters which displayed a densely interconnected map of early hematopoiesis in naked mole-rat BM (Figure 4A). Canonical cell cycle marker expression revealed cluster HSPC1 exclusively in G_1_ but not G_2_/M and S phase, whereas clusters ERY2/3 were not found in G_1_ (Figure 4B). Based on Uniform Manifold Approximation and Projection (UMAP) positioning we summarized closely connected groupings to collapse 16 clusters into 4 partitions. The HSPC1-4 partition contained stem cells and early lymphomyeloid progenitors as annotated by GSEA and consistent with the CD34 CITE-signal (Table S3, Figure 4C). The ERY1-5 partition was strongly associated with erythroid genesets while none of its clusters featured CD11b or Thy1.1 protein expression. To confirm our annotations and to profile expression kinetics across cells and clusters we highlighted differentially expressed genes with conserved roles during hematopoiesis (Orkin and Zon, 2008) (Table S4). In naked mole-rats GATA1 also determined erythroid commitment, as it was expressed from HSPC4 to ERY3, while GATA2 appeared to be expressed in subsets of HSPC1/2 and merged with GATA1 in HSPC4 and ERY1 (Figure 4D). Interestingly, GP9, expressed at the surface of platelets, was specifically produced in HSPC4 cells not expressing GATA2 (Figure S4H). This suggests HSPC4 as naked mole-rat MEPs with a GATA2-dependent switch between erythroblasts and megakaryocytic precursors. In HSPC1 HSCs HOXA9 expression dynamics created a corridor towards IL7R expression within HSPC3, marking the onset of lymphopoiesis in naked mole-rats (Figure 4E). Conversely, TARP expression displayed a gradient from HSPC2 to HSPC3 to be shut off in BC1, potentially portraying an early lymphoid progenitor (Figure S4G). We found 3 developmental stages of CD20^+^ BCs separated by gradual increase of CD79B (Figure S4F), with BC1/2 showing higher expression of VPREB1, located on proB and preB cells, than BC3 (Figure S4I). With the highest expression of mature BC markers CD79A/B (Figure S4F) and absence of VPREB1 in >90% of the cluster, BC3 maintained VPREB3 overexpression (Figure S4I), suggesting successive waves of VPREB genes as required for BC development in naked mole-rats. A small cluster (47 cells) likely comprised germinal center B or plasma cells (PC) as they highly expressed JCHAIN, MZB1 and EAF2 (Figure S4F). Another intriguing feature of naked mole-rat primitive hematopoiesis were expression patterns of ICAM2 vs EPCAM (Figure 4F): ICAM2, found on mouse fetal liver hemangioblasts (Pearson et al., 2010) and largely recapitulating the GATA2 pattern, depicted a path towards red lineage fate, while EPCAM marked lymphomyeloid commitment. (Notta et al., 2016) formulated a “two-tier” system with multipotency and MKs restricted to the stem cell compartment in adult humans, at the same time emphasizing that prenatally MK/ERY lineage branching occurs throughout the cellular hierarchy. Adult naked mole-rats seem to sustain oligopotent progenitors, in line with the ICAM2 expression trace from HSPC1 directly to HSPC4, which then splits into the erythroid route along GATA1^+^/GATA2^+^/EPOR^+^ vs the GATA1^+^/GATA2^−^/GP9^+^/CLEC1B^+^ axis towards MKs (Figure S4H). Likewise, EPCAM is induced in HSPC2 but peaks in HSPC3 (Table S4). Concomitantly, Thy1.1 decreased in equal fractions of both HSPC2/3, although CD11b was maintained (Figure 4C), pointing towards HSPC2 as a transient cell state between HSPC1 and lymphomyeloid commitment via HSPC2/3.

**Figure 4.**
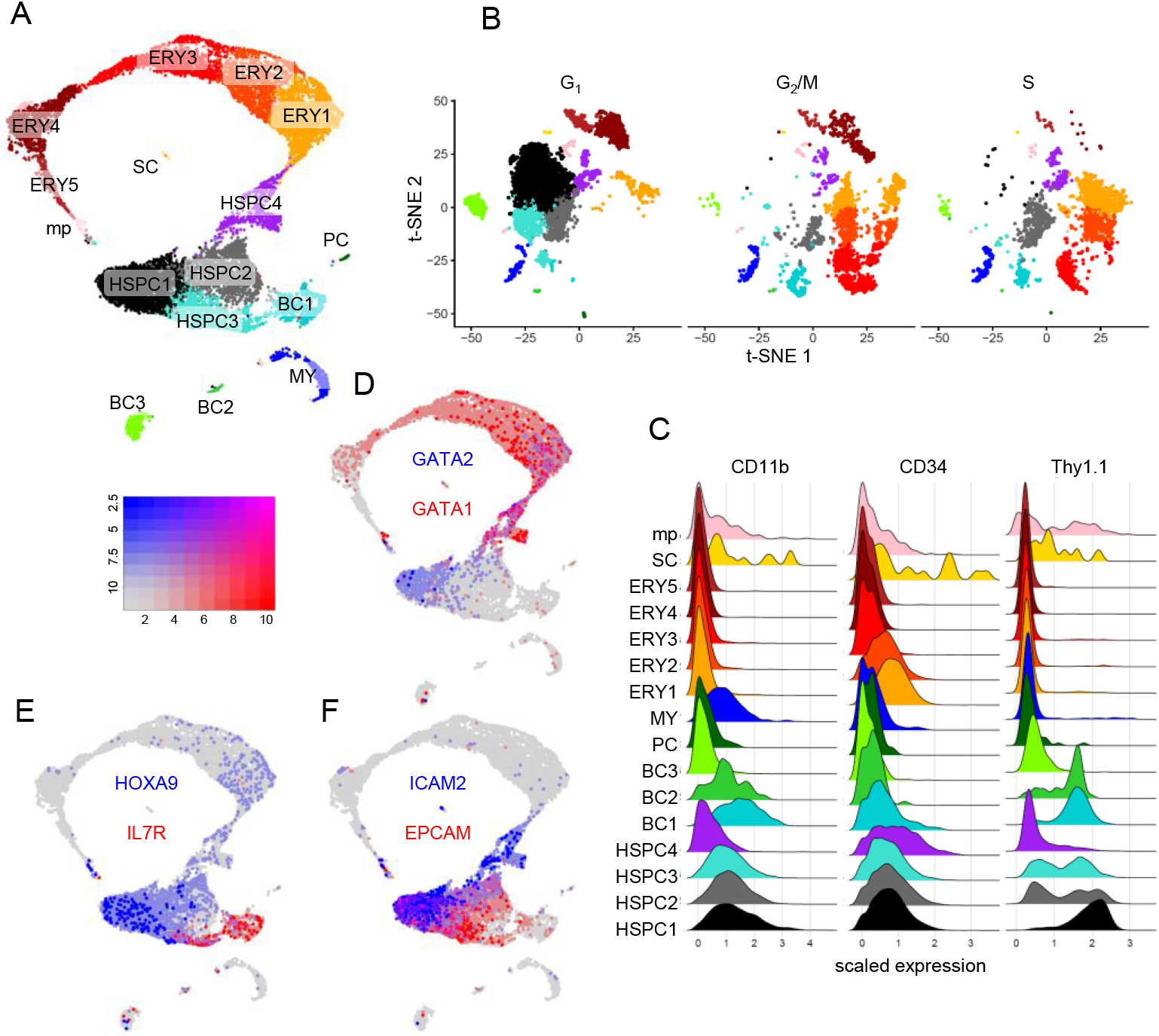
Single cell CITE-Seq of BM. (A) UMAP of 16 BM single cell clusters, 2000 most variable features, cell cycle regression. (B) T-SNE of 16 clusters grouped by canonical cell cycle marker expression. (C) CITE-Signals for each cluster, data centered and scaled. (D) Blendplot based on BM UMAP showing GATA1 (red, high expression), GATA2 (blue, high expression) and cells expressing both genes (purple). See scale on the left; expression, scaled UMI counts. (E) Blendplot showing BM IL7R vs HOXA9. (F) Blendplot showing BM EPCAM vs ICAM2. See also Figure S4.

Here we provide evidence for a bipotent MEP in adult naked mole-rats. This MEP stage appears uncoupled from the primitive leukocyte progenitor CP1 encompassing myeloid HSPC2 and lymphoid HSPC3 cells. The strict separation between red and white lineage priming resembles fetal linage choice and indicates retention of early hematopoietic stem cell programs in adult naked mole-rats. The stem cell state emerged as Thy1.1^+^/CD34^+^ cells expressing HOXA9, ICAM2 and CYTL1 (Figure S4G), a bone mass modulator exclusively induced in human CD34^+^ HSPCs (Shin et al., 2019). Strikingly, CITE-CD11b is expressed throughout the most primitive clusters HSPC1/2, and binary comparison of HSPC2 revealed that 14 of 26 genes upregulated vs HSPC1 are associated with mitotically active cells (Table S4-5). This supports the conclusion that the HSC state in naked mole-rats extends from LTCs into a fraction of dim LIN^+^ Thy1.1^++^/CD34^++^, which transition directly into lymphomyeloid-primed CP1 MPPs.

### TC development in naked mole-rats in and outside the thymus

Although we delineated B lineage priming by a primitive lymphoid progenitor in BM, the connection to TC differentiation remained elusive. We therefore profiled naked mole-rat thymi and found a LTC population (Figure 5A). A thymus-specific progenitor (TSP) fraction had Thy1.1^++^ levels identical to TCs and retained CD34^++^ levels of LTCs. There was a continuous transition from TSPs into a supposed early TC progenitor (ETP) state by successive loss of CD34, very similar to erythroid maturation (Figure 5A). A Thy1.1^+^/CD34^−^ population (CP6) at low frequency contains BC progenitors. We thus probed cervical naked mole-rat lymph nodes to determine their BC levels and found an inverted ratio of TCs and BCs as in thymi (Figure 5B, S4J-L). We noted that naked mole-rats have two types of cervical lymph nodes, one without LTC/TSP/ETP but high BC content (Figure 5B), and another with an immunophenotype identical to the naked mole-rat thymus (Figure 5C, S4L). Thymus and T lymph nodes share the same histological features such as many medullary sinuses, no germinal centers and more basophilic cytoplasms than B lymph nodes (Figure 5D-F), raising the possibility of extrathymic TC development under homeostatic conditions in naked mole-rats. Conversely, CD125^+^ BC frequencies are higher in B lymph nodes than in BM or spleens, suggesting those organs as the major BC site in naked mole-rats (Figure S4M-P).

**Figure 5.**
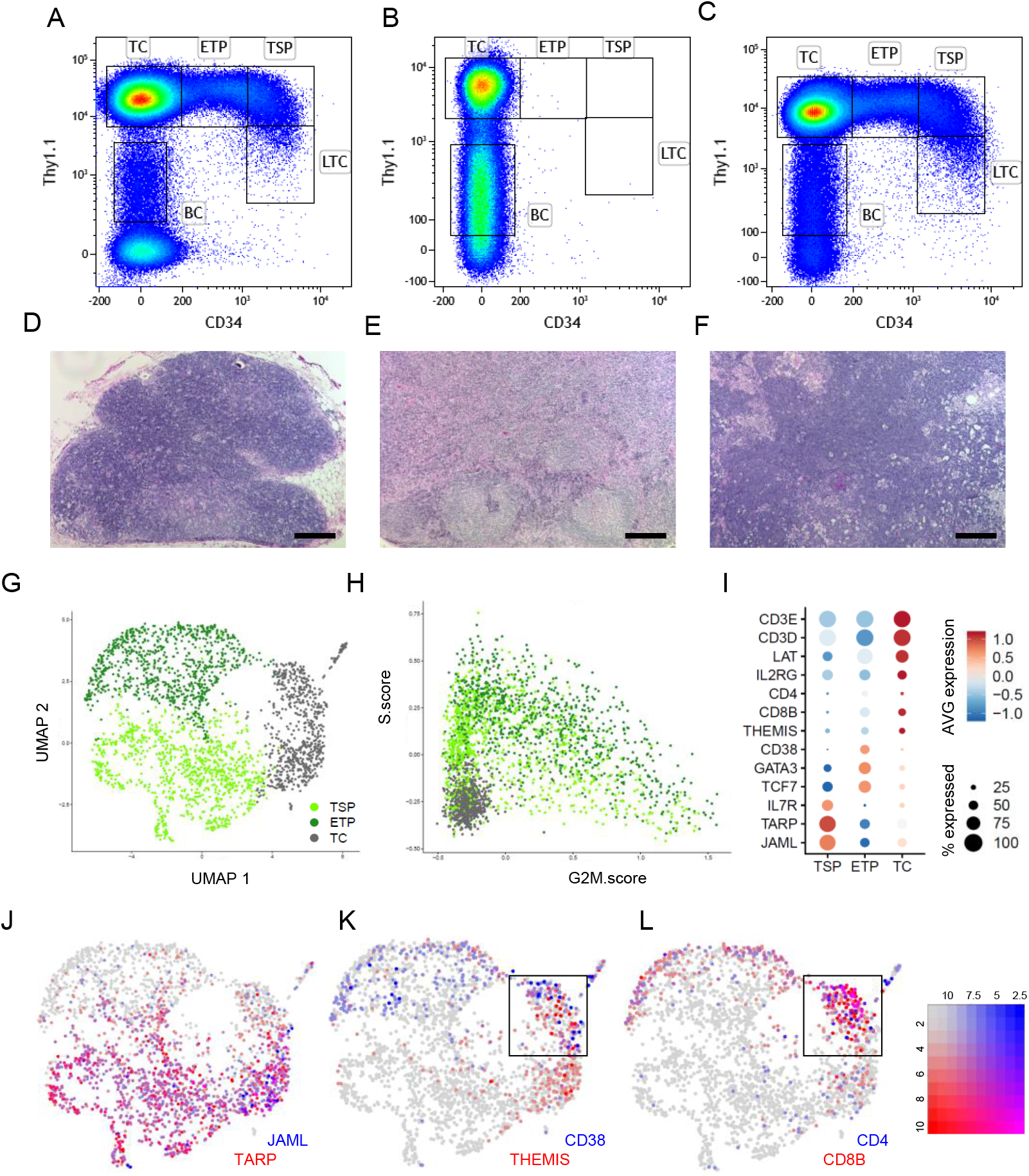
Lymphopoiesis in thymus and lymph nodes. FACS gating for Thymus (A), B-Lymph node (B) and T-Lymph node (C). LTC, LIN^−^ /Thy1.1^++^/CD34^++^; TSP, Thymus-specific progenitor; ETP, early T-cell progenitor; TC, T-cell; BC, B-cell. Hematoxylin & Eosin stained tissue sections of thymus (D), B-Lymph node (E) and T-Lymph node (F); Scale bars 200µm. (G) UMAP of 3 thymus single cell clusters, 2000 variable features, no cell cycle regression. (H) Cell cycle scoring of thymus clusters. (I) Dotplot showing expression of thymus marker genes; % expressed, cluster proportion with UMI ≥ 1. (J) Blendplot based on thymus UMAP showing TARP (red, high expression), JAML (blue, high expression) and cells expressing both genes (purple); see scale on the left. (K) Blendplot showing thymic THEMIS vs CD38. (L) Blendplot showing thymic CD8B vs CD4. See also Figure S4.

To better understand the thymic immunophenotype we performed CITE-Seq by sorting LTCs, TSPs and ETPs, yielding 8480 genes across 2466 cells with >1 mapped CITE label (Figure 5G, S4Q). Clustering based on gene expression separated 3 groups, which we annotated based on their CITE-label and cell cycle characteristics (Figure S4R). Cell cycle scoring revealed TSPs and ETPs as mitotically active progenitors (Figure 5H). The 3^rd^ cluster referred to TCs from the CD34^lo^ margin of the ETP sorting gate (Figure S4Q), since these cells expressed the highest levels of CD3E, LAT, CD247, CD4, CD8B and IL2RG while their S and G2M scores were <0 (Figure 4H-I). ETPs by contrast expressed CD4 and CD8B in larger fractions (33% and 57%, respectively vs 16% and 43% for TCs), and had higher levels of GATA3, an essential regulator of early TC development overexpressed at the DN3 to DN4 transition (Ho, I. C. et al., 2009). Unexpectedly, the Wnt gene expression program activating TCF7, which supports HSC self-renewal and diminishes CD34+ cell differentiation (Huls et al., 2013), was predominantly active in ETPs (Figure 5I). The axis IL7R-TARP was responsible for early lymphoid priming in naked mole-rat BM, and was retained in TSPs (Figure 5I). Interestingly, JAML has been shown to induce γδTC activation (Witherden et al., 2010) and is stringently co-expressed with TARP (Figure 5J). CD38, which is expressed mainly in thymocytes at the double-positive stage (Tenca et al., 2003), and THEMIS expression, indispensable for proper positive and negative TC selection in mice (Johnson et al., 2009), converged in a cell cluster enriched with CD4 and CD8B expressing cells (Figure 5K-L). Therefore, within this cell cluster positive and negative αβTC selection occurs. NOTCH1 directs αβTC development during β-selection, whereas ID3 restricts thymic γδTC development (Lauritsen et al., 2009). Conversely, in the naked mole-rat thymus ID3 was expressed in the cell cluster of TC selection and in ETPs, while NOTCH1/TARP/JAML expression marks a cell state transition from TSPs towards αβTC development (Figure S4S).

In summary we provide evidence for extrathymic TC development in healthy naked mole-rats. Thymic involution occurs in almost all vertebrates (Shanley et al., 2009), hence the presence of T lymph nodes in the naked mole rat may serve as a way to compensate for thymic involution and maintain youthful TC-mediated immune function during adulthood. TARP as a transcription product of an invariant TC receptor locus emerged as a central player during TC development unique to naked mole-rats. TARP may instruct γδTC fate, which could explain the low frequency of CD4 and CD8 TCs in blood and would propose high frequencies of CD4^−^CD8^−^ γδTCs in naked mole-rats.

### Pseudotemporal ordering remodels naked mole-rat hematopoiesis

The dynamic process of differentiation and organogenesis can be modeled using trajectory inference methods (Saelens et al., 2019). We combined the scRNA-Seq datasets from blood, BM and thymus to a joint dataset comprising 18054 cells and 10441 orthologues divided into 19 clusters. Developmental annotation retrieved human HSC and naked mole-rat LTC signatures for the 3533 cell HSC cluster, comprising 19.6% of all sequenced cells and converging with 17.4% sorted CP1/2 cells (Figure S5A-B, Table S6). Dimensionality reduction displayed a structure of the HSC/MPP/BCP/MEP cluster node analogous to the BM HSPC partition, whereas most common cell type overlaps across tissues were observed with mature BCs, TCs and DCs (Figure S5C-D). Differential expression analysis found 2165 unique genes specifically upregulated in one or more clusters (Figure 6A, Table S4). We applied partition-based graph abstraction (PAGA) (Wolf et al., 2019) to capture UMAP connectivity features (Figure 6B). The trajectory rooted in the HSC/MPP clusters and trifurcated into erythroid, myeloid and lymphoid paths. A stem cell state was portrayed by pervasive expression of TM4SF1 (Figure 6C): this is the top HSC marker by fold-change and differential detection ratio, also a marker for CP1/2, amongst the gradually declining genes from CP2 to CP1, and the top BM HSPC1 marker (Table S2, S4). At single-cell resolution TM4SF1 increased in one subset of MPPs, while absent in 2 others. Specifically, MPPs appeared to trifurcate into a stage with high TM4SF1/EPCAM (Figure S5E), TM6SF1/CEBPα/CEBPε towards myeloid progenitors (BM.MY; Figure 6D, S5F-G) and via IL7R/TARP towards lymphoid commitment (BCP, Figure S5H-I). Displaying the trajectory milestone connectivities along pseudotime pictured a root stem cell state, which linked to the nearest neighbor cluster MPP (Figure 6E). The dendrogram fortified a direct HSC/MEP transition and precisely recapitulated CD34 FACS and BM CITE-Seq clustering patterns along the erythroid lineage. We confirmed our data using a minimal-spanning tree (Qiu et al., 2011), which found a lymphomyeloid branching point between two HSC states and highlighted the direct HSC/MEP connection towards erythroid commitment as a separate milestone (Figure S5J-K).

**Figure 6.**
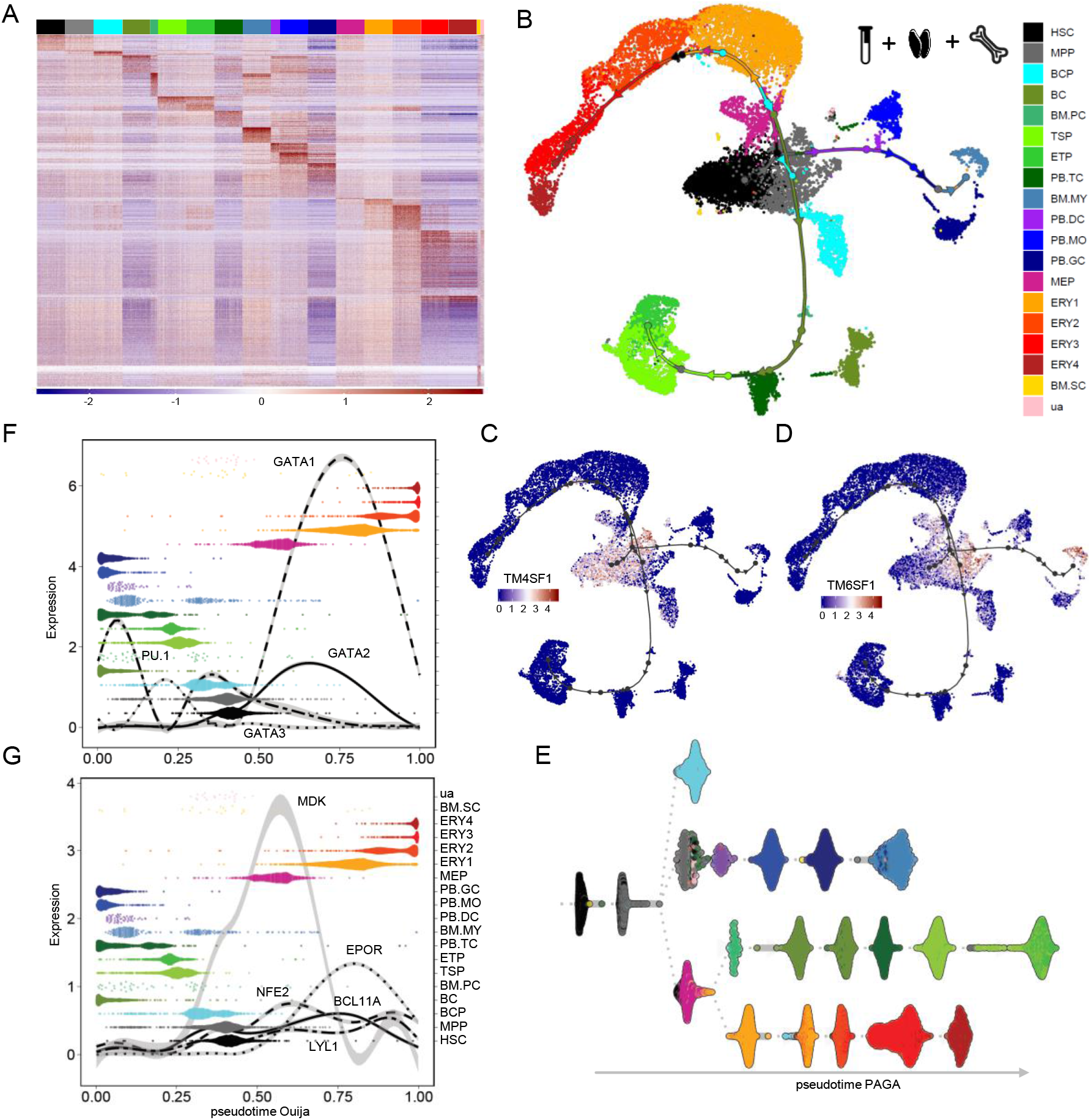
Pseudotemporal ordering of the hematopoietic hierarchy. (A) Heatmap showing 2165 overexpressed marker genes of 19 single cell clusters from BM, blood and thymus randomly downsampled to `≤` 200 cells; color coding is used throughout this Figure. (B) PAGA cell trajectories plotted over UMAP dimensionality reduction, see legend for color code. Connecting line, trajectory; Circles, trajectory milestones; Arrowheads, pseudotime direction; coloring by nearest cell. UMAP highlighting TM4SF1 (C) or TM6SF1 (D) expression throughout the trajectory; Expression, log2 UMI counts. (E) Dendrogram of PAGA pseudotime with Louvain clustering input, root set to a random seed HSC cluster cell. (F, G) *Ouija* pseudotime (x-axis) for indicated gene expression (left y-axis), cells ordered into Louvain cluster categorials (right y-axis); Expression curves smoothed using GAM models. See also Figure S5-6.

Aiming to provide an ordering adapted to a multibranched hierarchy we applied the supervised *Ouija* pseudotime inference (Campbell and Yau, 2019) using 28 switch and 23 transient genes curated from differential gene expression analysis. The GATA family congruently remodeled hematopoietic trajectories along Ouija pseudotime: GATA1 was strictly confined to erythropoiesis and sharply increased at the boundary of HSC/MEP, high GATA3 levels were limited to ETPs. Low GATA2 abundance marked the stem cell state, from which it increased to follow GATA1 throughout erythroid fate (Figure 6F). PU.1 is a transcription factor of the myeloid lineage with a role in HSC maintenance (Staber et al., 2013). Likewise its expression was highest in BM.MY and PB.DCs, and while PU.1 was induced in naked-mole rat MPPs, its expression declining in MEPs suggests conservation of the classic GATA1-PU.1 bi-modal switch (Figure 6F). Diffusion maps, applying unsupervised random walk paths (Angerer et al., 2016), also yielded a central position of the naked mole-rat stem cell compartment, albeit with less sharp cluster distinction (Figure S5L). We found that BCL11A had a role in naked mole-rat erythropoiesis, as its levels increased over MEPs and ERY1, similar to EPOR expression patterns (Figure 6G). Interestingly, both NFE2 and LYL1, known erythroid fate regulators, followed a biphasic trend with peaks at MEP and ERY2 stage. The pleiotropic neurite growth factor midkine emerged as a specific marker of naked mole-rat MEPs (Figure 6G, Table S4).

In summary, pseudotemporal ordering of a cytometry-enriched naked mole-rat HSPC compartment defined HSCs, MPPs, confirmed the HSC/MEP-transition and allowed mapping the principal myeloid and lymphoid trajectories through a common primitive progenitor. The presence of a distinct MEP may be beneficial to longevity by restricting the path towards myeloid differentiation and counteracting age-related myeloid bias.

### Comparative biology reveals effective mitochondrial utilization in naked mole-rat HSPCs

With a primary map of the naked mole-rat hematopoietic system we directly compared RNA-Seq transcriptomes between HSPCs of 3 species, encompassing 9062 orthologues across 59 samples from 10 distinct cell states (Figure S6A). The highest separation index was interspecies clustering, whereas intraspecies cell type associations recapitulated single species t-SNEs (Figure 7A). A total of 7194 variable features between the three species were fed into ssGSEA using the hallmark genesets, yielding 28 top hit pathways (Figure 7B, Table S5). In this comparison, terms HYPOXIA, GLYCOLYSIS and APOPTOSIS had highest enrichment scores for human cells. In murine cells the 3 top terms were G2M_CHECKPOINT, MITOTIC_SPINDLE and E2F_TARGETS, while MYC_TARGETS_V1 ranked with most positive enrichment scores, Supporting elevated cycling and metabolic activities of mouse BM. Interestingly, both human and mouse enriched for PI3K_AKT_MTOR_SIGNALING, while naked mole-rat cells showed relation to MTORC1_SIGNALING. Naked mole rat HSPCs showed increased OXPHOS and ADIPOGENESIS, and reduced INFLAMMATORY SIGNALLING and IL2_STAT5 signaling. Decreased inflammatory signaling in HSPCs may reflect a less inflamed stem cell niche in the bone marrow.

**Figure 7.**
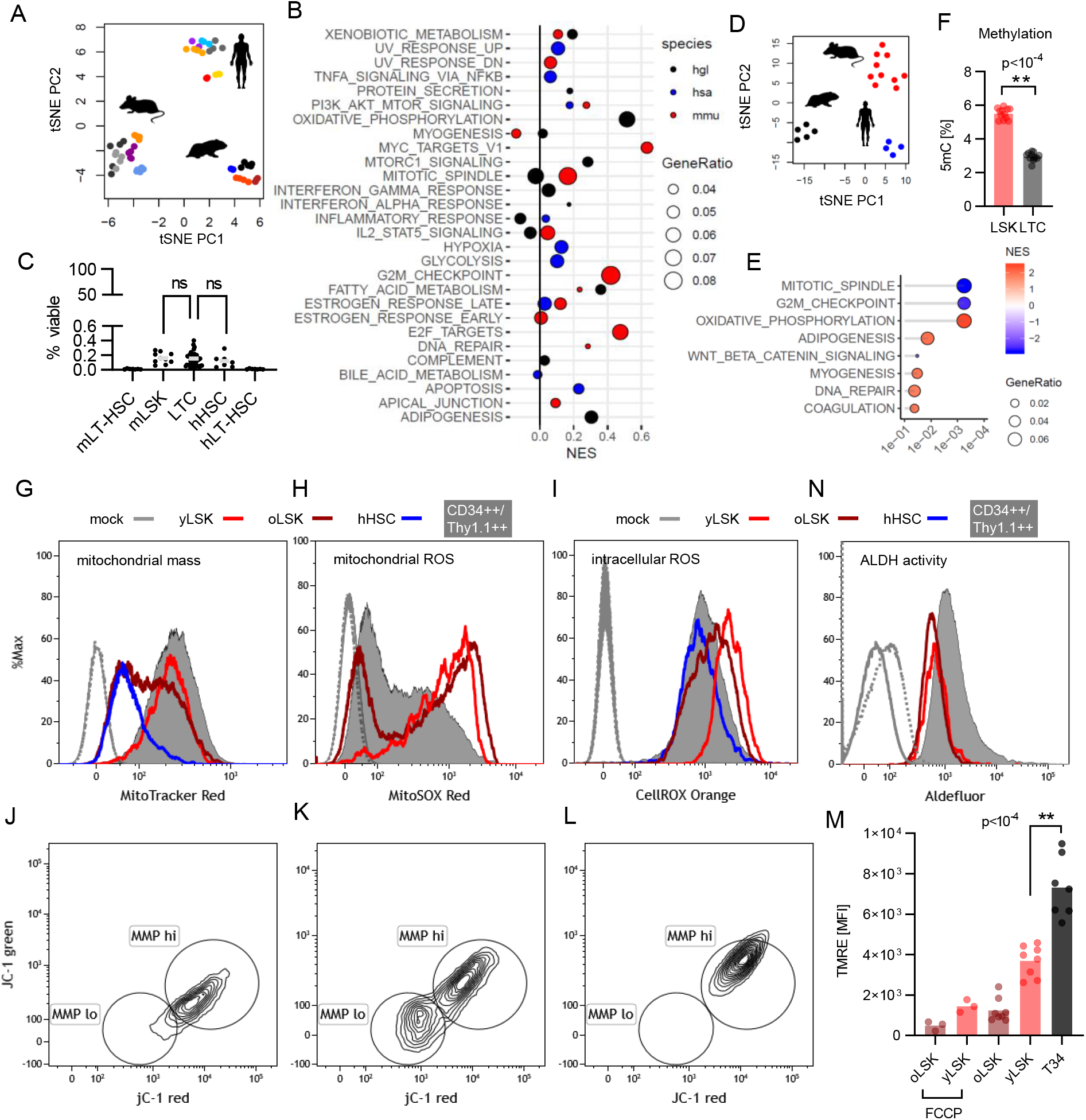
Comparative transcriptomics reveal elevated OXPHOS activity in naked mole-rat HSPCs. (A) Unsupervised t-SNE clustering, color code see Figure used on heatmaps this Figure 2E, S2I, S2N. (B) ssGSEA of HSPCs displaying the top 28 terms filtered for q < 10^-3^ and GeneRatio > 0.03. hgl, naked mole-rat; hsa, human; mmu, mouse; color code used throughout the figure. (C) Frequencies of BM HSC compartments between species. mLT-HSC (N=7), mouse LIN^−^/Sca-1^+^/Kit^+^/CD48^−^/CD150^+^; LSK (N=10), mouse LIN^−^/Sca-1^+^/Kit^+^; LTC (N=47), naked mole-rat LIN^−^ /Thy1.1^+^/CD34^+^; hHSC (N=7), human LIN^−^/CD38^lo^/CD34^+^; hLT-HSC (N=7), human LIN^−^/CD38^lo^/CD34^+^CD45RA^−^/CD90^+^; p-value determined by Dunnett’s One-way ANOVA. (D) Unsupervised t-SNE clustering of mouse LT-HSCs & ST-HSCs, human CB-HSCs and naked mole-rat LTCs across all conserved orthologs. (E) fGSEA of LTC 3-species signature displaying the top 8 terms filtered for q < 0.05. (F) Global DNA 5-Methylcytosine (5mC) levels. p-values were determined by unpaired Welch’s t-test; N=12. Signal Intensities for MitoTracker (G), MitoSOX (H), CellROX (I) and Aldefluor (N); % max scaling used on each histogram. Samples from each group were merged; Continuous grey curves, mouse unstained (mock); Dotted grey curves, naked mole-rat unstained (mock). Merged JC-1 fluorescence biplots for young LSK (yLSK; J), old LSK (oLSK; K), naked mole-rat Thy1.1^++^/CD34^++^ (L). (M) Mean fluorescence intensities of TMRE stainings. p-values were determined by Tukey’s One-way ANOVA; LSK, N=8; T34 (Thy1.1^++^/CD34^++^), N=7. See also Figure S6-7.

Remarkably we found no significant differences in the frequencies of LTCs, LSKs and hHSCs in normal BM (Figure 7C, S6B-C), suggesting conserved stem cell pool frequencies. Thus, we compared HSCs from the 3 species to find 1415 up- and 1789 downregulated genes specific for LTCs (Figure 7D). A notable finding was the strong negative association with cell cycle terms (Figure 7E, Table S5). Indeed, Ki67 staining showed a higher cell fraction in G_0_ for LTCs as compared to LSKs (Figure S6D). It is interesting to note that mouse LKs containing CMP/GMP/MEP had the same cycling properties as CP1 comprised of HSC/MPP/BCP (Figure S6D). One of the downregulated genes in LTCs vs other species HSCs was DNMT1 (38.5-fold, p<10^-6^), which is essential for HSC self-renewal (Broske et al., 2009). DNMT1 levels were repressed in primitive LTCs compared to CP1 cells (Figure S6E). Strikingly, total DNA 5-Methylcytosine (5mC) levels are reduced by 1.9-fold in LTCs vs mouse LSKs (Figure 7F).

The strongest enrichment in naked mole-rat HSPCs was OXPHOS (Figure 7B-E). Mapping all OXPHOS signature genes to the respective KEGG pathway depicted all five complexes of the respiratory chain each with at least one overexpressed component (Table S7). As we did not detect any differences between naked mole-rat LTCs and CP1 in any of the following measurements, we gated on viable Thy1.1^++^/CD34^++^ (T34; Figure S6F), which have different mitochondrial loads than TCs or erythroid cells (Figure S6G). Next we probed mitochondrial mass of HSCs from 3 species to find no significant differences between naked mole-rat and mouse old (oLSK, 25 months) and young (yLSK, 3-4 month) HSCs (Figure 7G, S6H). However, human HSCs have fewer mitochondria than their rodent counterparts. Respiratory activity generates ROS especially in mitochondria, therefore we quantified mitochondrial superoxide levels between HSCs (Figure 7H). Although a trend progressed towards less mitochondria in oLSK vs yLSK and there was higher staining in yLSK than oLSK, we found much less superoxide in naked mole-rat T34s (Figure S6I). Intracellular ROS levels showed more ROS for mouse yLSKs than for T34s and human HSCs (Figure 7I, S6J). The mitochondrial membrane potential (MMP) as a resultant of OXPHOS is an indicator of mitochondrial activity. In line with (Ho, T. T. et al., 2017) we showed that oLSKs feature an increased fraction of cells with low MMP compared to yLSKs (Figure 7J-L, S6K). Old HSCs increase frequency through higher cell cycle activity (Kowalczyk et al., 2015), thereby relying on glycolytic metabolism, hence oLSK ROS levels are decreased compared to yLSKs with higher OXPHOS activity. In this regards an intriguing finding is that naked-mole rat HSCs have even higher OXPHOS than both human and mouse while showing less ROS and superoxide levels, pointing towards higher efficiency in ROS clearance or avoidance. Moreover, T34 cells have no low MMP fraction and increased fluorescence levels of JC-1 red, indicating a higher basal MMP than murine cells (Figure 7L). JC-1 red is even higher in ERY cells, which we showed to harbor more mitochondria than T34s (Figure S6L). We confirmed the higher MMP in T34s compared to yLSKs using Tetramethylrhodamine (TMRE) sequestration (Figure 7M).

The β-oxidation of fatty acids provides substrates for OXPHOS, and naked mole-rat HSPC gene expression was enriched for terms ADIPOGENESIS and FATTY_ACID_METABOLISM. Aldehyde dehydrogenases (ALDH) metabolize aldehydes arising from processes such as lipid peroxidation. Also, an ALDH “bright” subset of CD34^+^ CB-HSCs has been reported to enrich in HSCs (Storms et al., 2005). ALDH staining revealed 2-fold higher levels in T34s compared to yLSKs (Figure 7N, S6M), another cytoprotective feature upregulated in naked mole-rat cells.

As a classical marker in HSC research Rhodamine 123 (Rho) efflux has been shown to functionally enrich human HSCs by low Rho levels and was used to purify mouse HSCs obtained by the side population (SP) phenotype (Uchida et al., 1996; Uchida et al., 2003). Lack of CD34^+^ expression in SP cells from human CB has been reported earlier (Pearce and Bonnet, 2007), likewise we did not detect a reliable SP in naked mole-rat BM (Figure S7A-B). Consistent with an enrichment in genes associated with XENOBIOTIC_METABOLISM we detected a stronger Rho efflux in T34s than in LSKs (Figure S6N-O).

Taken together, these data suggest that naked mole-rat HSCs have evolved a mechanism of stem cell homeostasis involving elevated OXPHOS and improved clearance of damaging compounds. OXPHOS is more energy-efficient and prevents Lactate-caused acidification, which likely contributes to preservation of quiescence and tissue homeostasis during aging.

## DISCUSSION

Naked mole rats are the longest-lived rodents, and remarkably they remain healthy until the end of their lives and are resistant to age-related diseases including cancer. As stem cells are essential for maintenance and repair of tissues, stem cell biology of the naked mole rat is of immediate interest to biomedical research.

Here we present a comprehensive analysis of the blood system in >60 naked mole-rats including functional and molecular characterization of stem and progenitor subtypes and a primary landscape of the hematopoietic hierarchy. We identified several unique features that may contribute to longevity of the naked mole rats. Some of these features, such as splenic hematopoiesis, are characteristic of embryonic stage in other mammals. It had been proposed that long-lived naked mole rats, as well as humans, display neotenic traits compared to their short-lived relatives (Skulachev et al., 2017). Furthermore, many characteristics of the naked mole rat hematopoietic system showed higher similarity to humans than to mice (Figure S7D).

A striking feature of naked mole rat stem cells was a higher proportion of cells in quiescence. The dynamic equilibrium between quiescence and specific cell cycle kinetics is a hallmark of adult stem cells, hence mouse HSCs have been shown to contain a dormant fraction of ∼20% (Foudi et al., 2009). Notably, CDK6 expression is a quiescence marker for human HSCs (Laurenti et al., 2015), whose levels are identical between hHSCs and LTCs (Figure S6E). Dormancy is also associated with high levels of CDKN1C (Zou et al., 2011), which is most abundant in LTCs compared to human and mouse HSCs (Figure S6E). Altogether accumulating evidence suggests an expanded quiescent HSC pool in naked mole-rats (Figure S6D) and rigid control of cell cycle genes at the transcriptional level (Figure 3B, 4B, 7E). The quiescent HSC pool would benefit longevity by minimizing damage to stem cells and preventing clonal expansion, which is a key feature of an aged hematopoietic system (Zink et al., 2017).

The HSC state features specific stress response and quality-control mechanisms to preserve its integrity after exposure to DNA damage or metabolic stress. Metabolic paradigms of HSCs are their reliance on glycolysis and low mitochondrial activity (Simsek et al., 2010; Vannini et al., 2016). By contrast, naked mole-rats had an overexpressed OXPHOS signature and elevated MMP, but similar HIF1A levels than human HSCs (Figure S6E). Importantly, naked mole-rat HSPCs expressed key enzymes of the HIF1α-mediated glycolytic switch (PDKs) as wells as glycolytic enzymes (HK, PFK) and IDH homologs at higher degrees than human and mouse cells (Figure S7C), pointing towards increased glycolytic activity for subsequent OXPHOS substrate production. Notably, human HSCs naturally live longer than their rodent counterparts, have fewer mitochondria but produce the same amount of ROS as naked mole-rat HSCs, suggesting robust antioxidant responses as a key mechanism to preserve stemness in the naked mole-rat. Additionally, the glycolytic state is a key characteristic of cancer cells, which makes human and mouse HSCs one step closer to oncogenic transformation. Hence, overexpressed OXPHOS in the naked mole rat HSCs may be protecting them from hematopoietic malignancies.

A common hallmark of hematopoietic lineage trajectories is a differentiation bias of HSCs along the MK lineage (Sanjuan-Pla et al., 2013). However, this bias is shifted in naked mole-rats towards HSC/MEP transition, resembling the fetal HSC bias towards the entire red lineage, therewith constituting a further neotenic trait of naked mole-rats (Skulachev et al., 2017). Concordantly, adult MKs and platelet levels are reduced compared to mice, and LTCs and CP1 cells expressed MPL at lower levels than mouse and human HSCs and CMPs (Figure S6E, S7D). This likely contributes to reduced fatal thrombotic in old age.

A very striking finding was the presence of two types of lymph nodes in the naked mole-rat: B lymph nodes containing germinal centers and maturing B cells, and T lymph nodes showing similar histology to thymus and T cell progenitors. Thymic involution occurs in the majority of vertebrates (Shanley et al., 2009), resulting in decreased output of naïve T cells and reduced ability to mount protective responses against new antigens. The presence of T lymph nodes in the naked mole-rat may serve to overcome this age-related change and prevent immune-senescence. It would be interesting to investigate whether T lymph nodes maintain their function with age, however, as we do not have animals older than 10 years of age in our colony we could not address this question.

Shared transcription factor networks between mouse HSCs and LMPPs impose a BC differentiation trajectory in the latter (Laurenti et al., 2013). This pattern surfaced in naked mole-rat HSPC4/BCP cell states, which also express TSP markers IL7R and TARP, hence likely qualify as multilymphoid progenitors (MLP). Thymic TARP/JAML co-expression and an abundant CD4^−^ CD8^−^ cell fraction in blood raise the possibility that invariant TC receptor cells comprise the majority of the TC pool. While αβTCs primarily function in foreign antigen recognition, γδTCs mediate tissue homeostasis and damaged self recognition (Nielsen et al., 2017; Silva-Santos et al., 2019). The shift from αβTC-towards γδTC-mediated immunity may provide means to sustain tissue integrity and exert enhanced anti-tumor functions throughout their long lifespan.

In conclusion, naked mole-rat stem cells appear more quiescent than their murine counterparts, feature elevated mitochondrial function and lower ROS levels. Overall, the entire hematopoietic system of naked mole-rats evolved a combination of unique adaptations to an extended healthspan, such as diminished platelets and megakaryocytes, active hematopoiesis in the spleen and extrathymic TC development in T lymph nodes. The naked mole-rat has evolved extreme longevity and resistance to almost all age-related diseases. Understanding the molecular mechanisms of these evolutionary adaptations can lead to novel strategies to improve human health. Our resource provides a platform for using naked mole-rats as a research model in stem cell biology, immunology, inflammation and the studies of systemic factors in aging.

## Supporting information

Table S1 FRAMA transcripts

Table S2 population RNA-Seq signatures

Table S3 scRNA-Seq BM partitions GSEA

Table S4 scRNA-Seq signatures

Table S5 genesets

Table S6 BM PB THY clusters GSEA

Table S7 KEGG pathway OXPHOS

## STAR METHODS

Detailed methods are provided in the online version of this paper and include the following:

- CONTACT FOR REAGENT AND RESOURCE SHARING
- EXPERIMENTAL MODEL AND SUBJECT DETAILS
  - Animals
  - Primary cell isolation
- METHOD DETAILS
  - Hematology Analyzer
  - Histology
  - Global methylation levels
  - Total BM cellularity
  - Methylcellulose Colony Assays
  - Flow Cytometry
  - Xenotransplantations
  - Transcriptome Assembly
  - Population RNA-Sequencing
  - Single-cell RNA-Sequencing
- QUANTIFICATION AND STATISTICAL ANALYSIS

## SUPPLEMENTAL INFORMATION

Supplemental Information can be found in the accompanying Manuscript_Supplement.pdf file.

## ACKNOWLEDGMENTS

Many thanks to Alex Aiezza II for RSEM builds of mouse samples, Anthony Corbett for FRAMA and Jason R Myers for a FRAMA-into-RSEM interface and cellranger processing of scRNA-Seq data. Gratitude goes to Leonid Peshkin and helpful hints from SPRING. BM from healthy human donors were a gift from Prof Liesveldt. Special thanks to Jan-Henning Klusmann for critical comments on the manuscript. This work was supported by the US National Institutes of Health grants to V.G. and A.S. S.E. is a fellow of HFSP.

## AUTHOR CONTRIBUTIONS

S.E. designed and supervised research, performed most experiments and analyzed data; M.M. performed FRAMA-genome alignments; M.E.S. performed histology and most histochemistry stainings; A.T. assisted with FACS; M.T. and V.N.G. contributed to data analysis; A.S. and V.G. supervised research; S.E., A.S. and V.G. wrote the manuscript with input from all authors.

---

**Figure S1.**
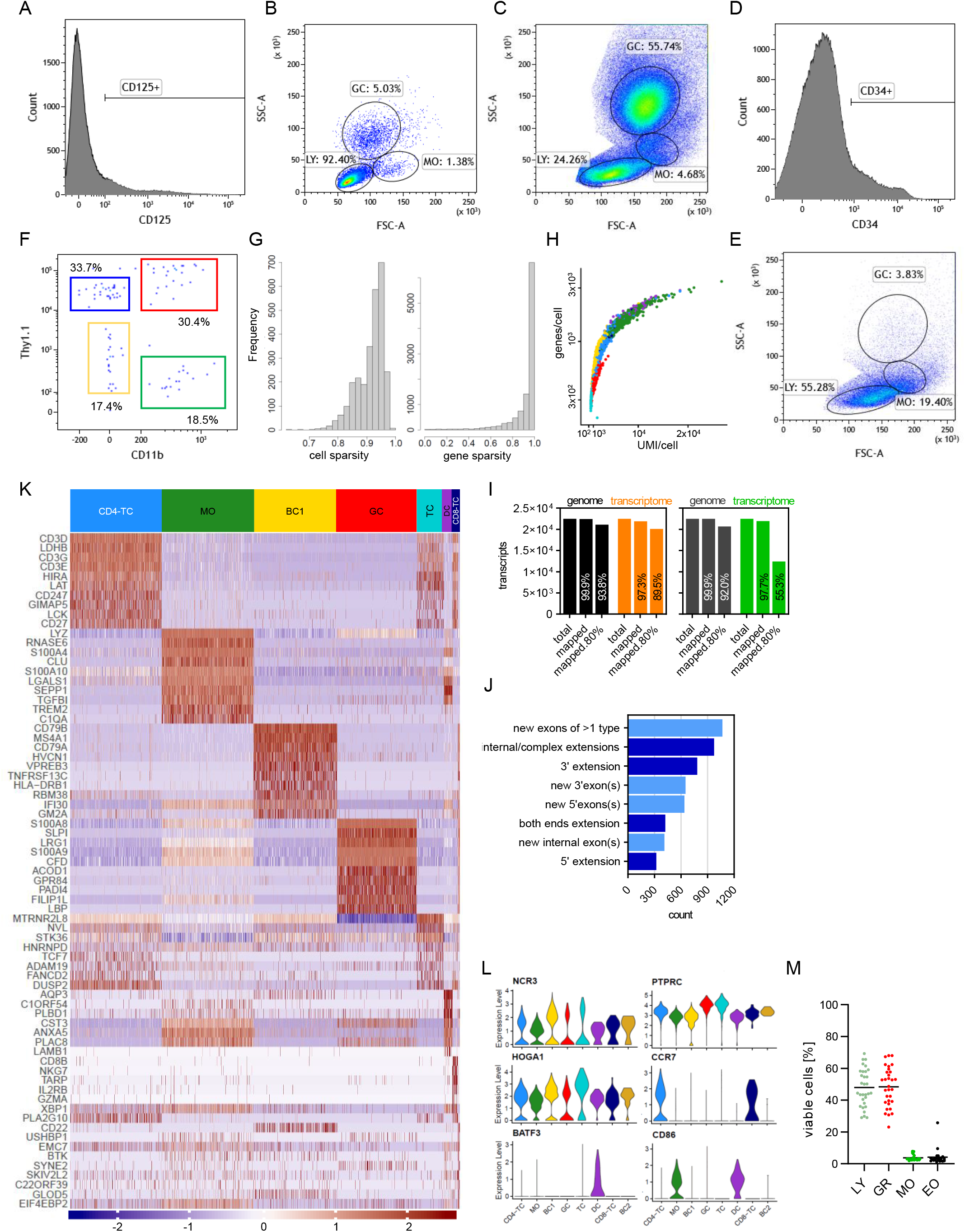

**Figure S2.**
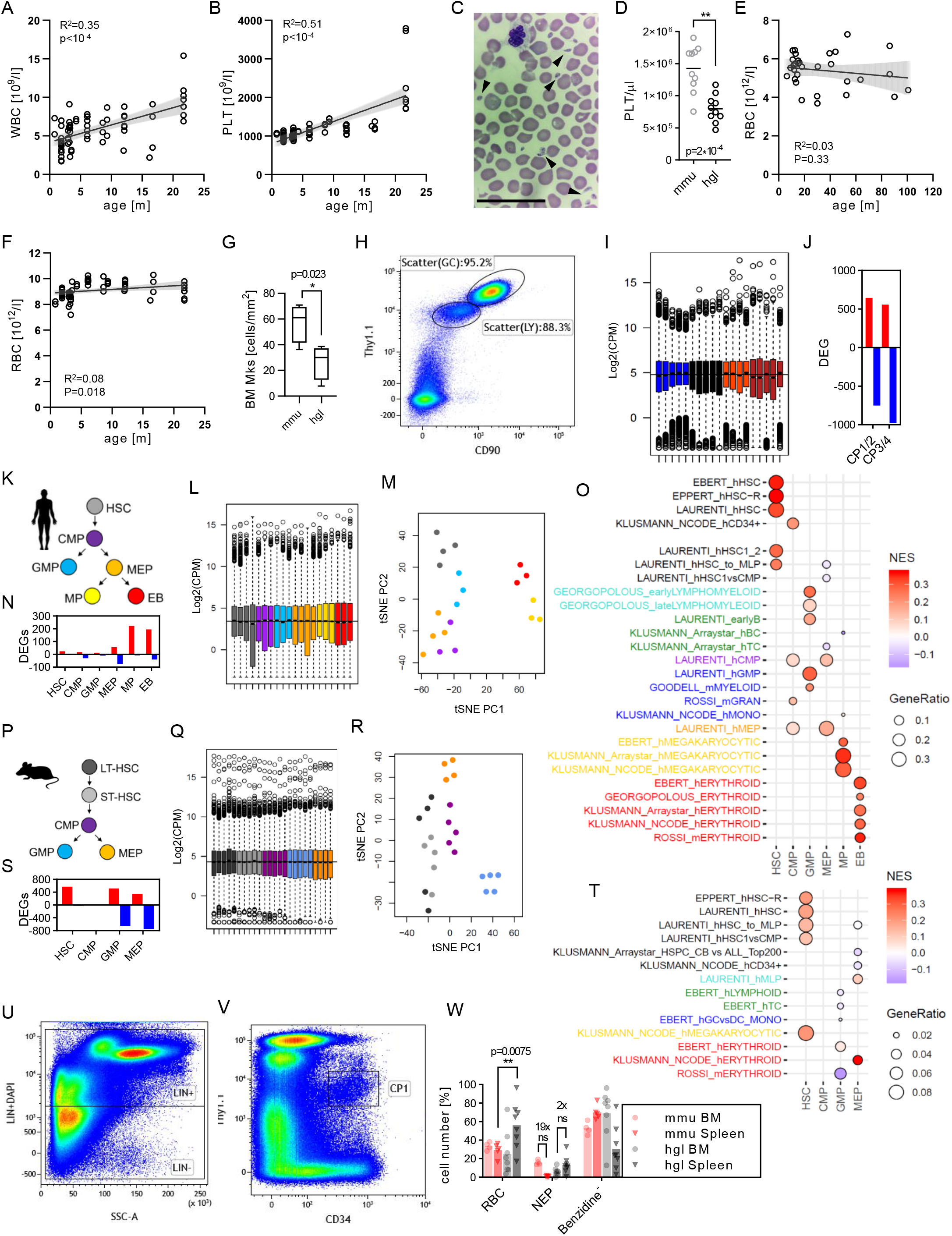

**Figure S3.**
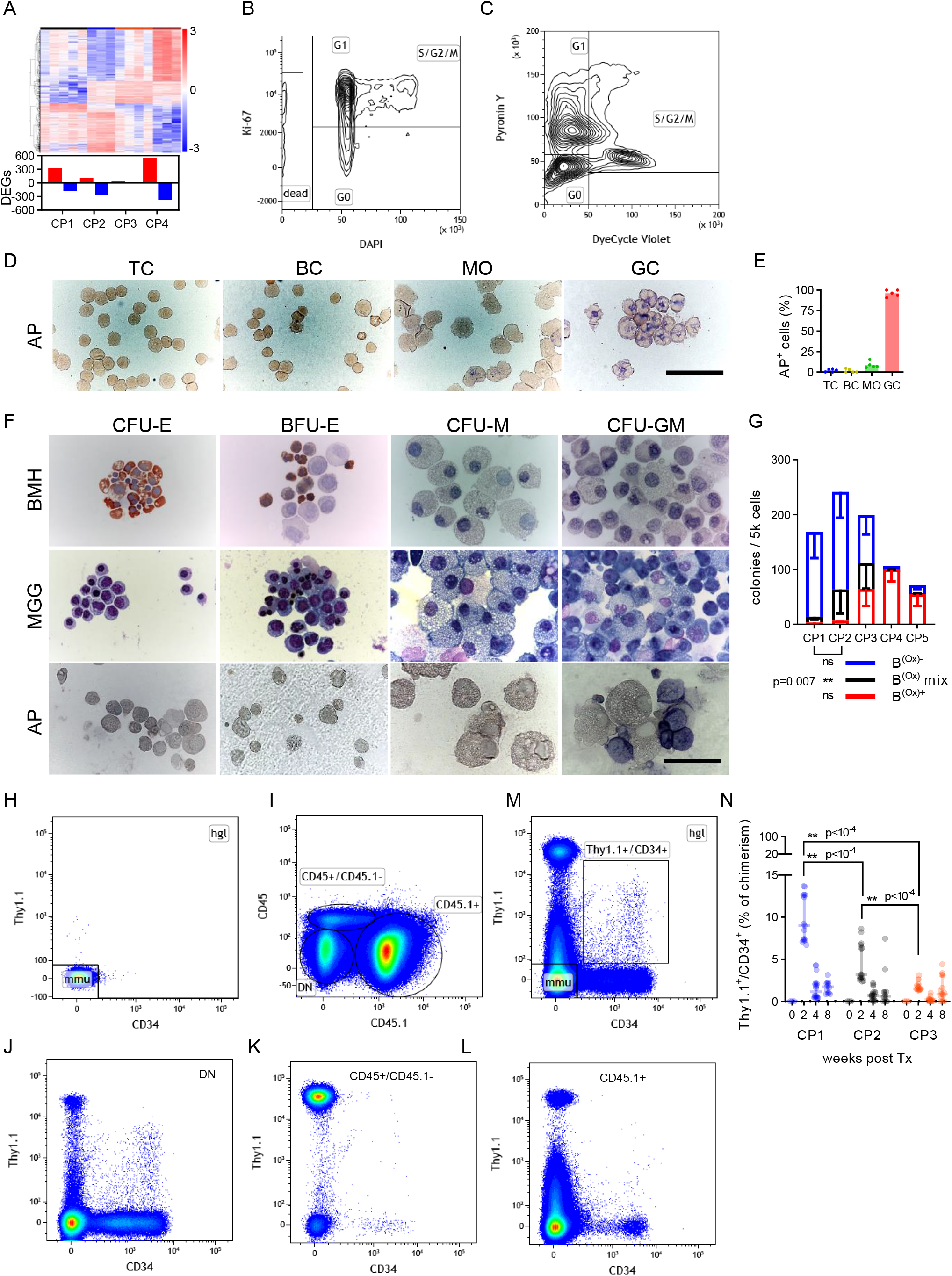

**Figure S4.**
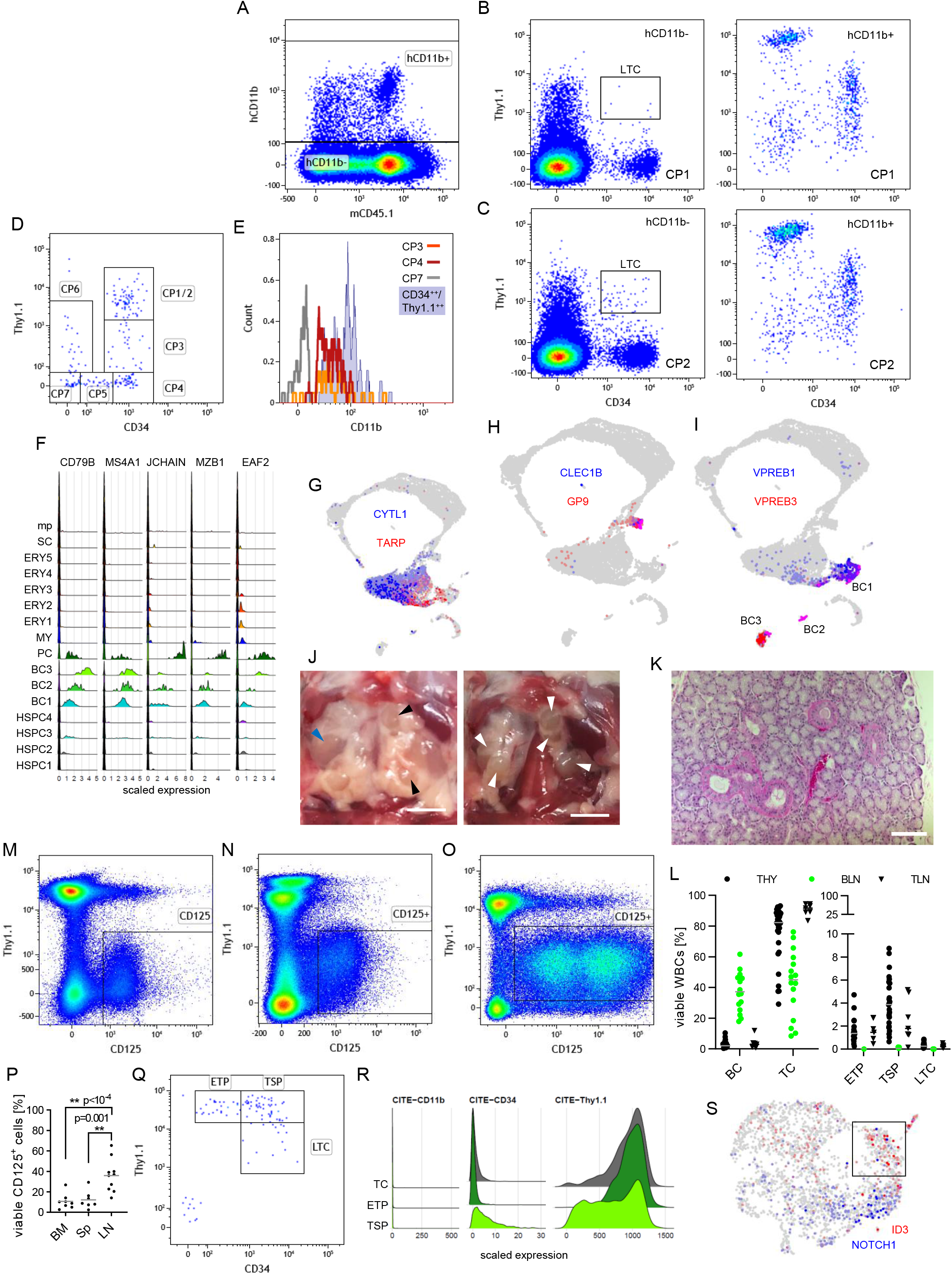

**Figure S5.**
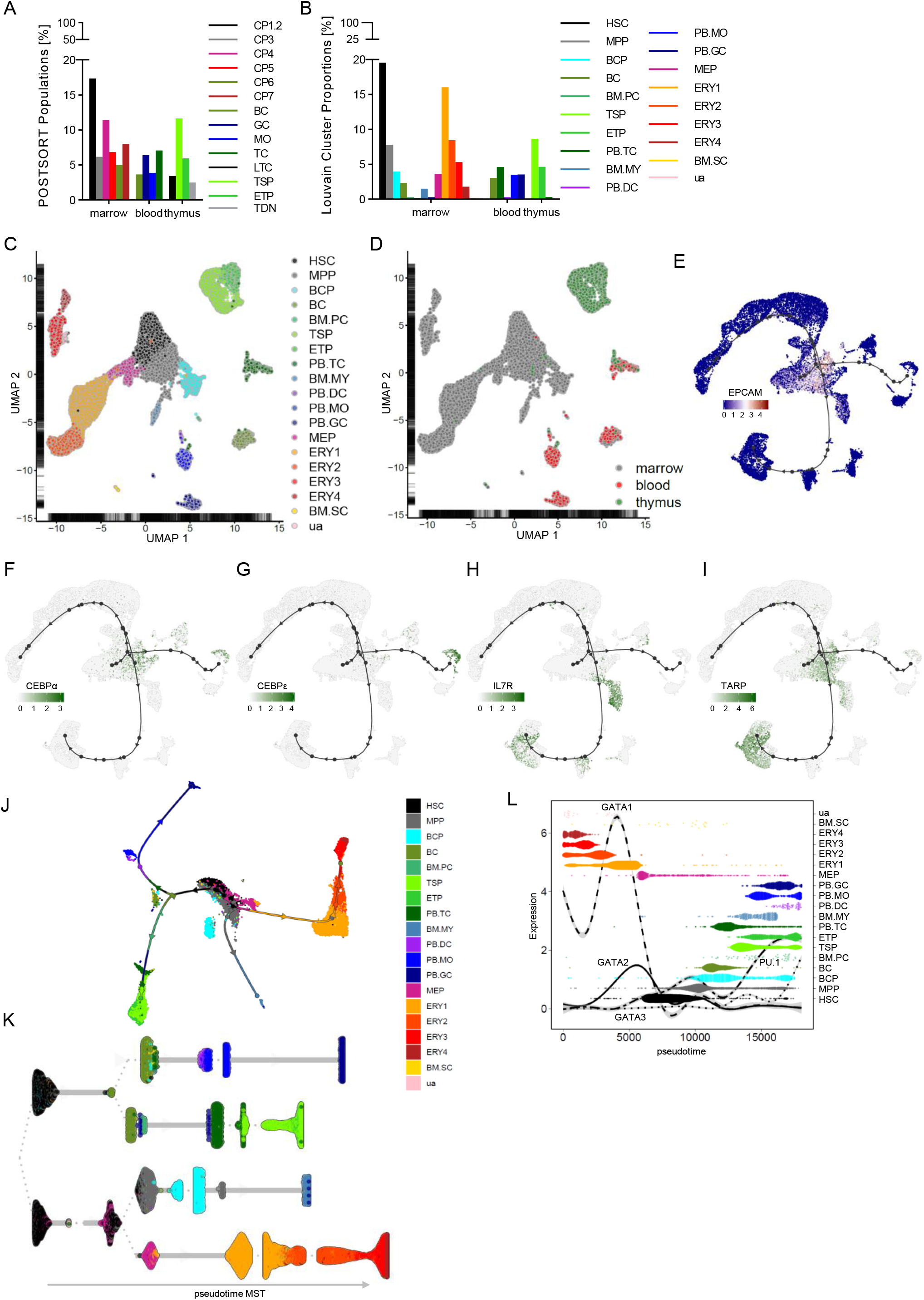

**Figure S6.**
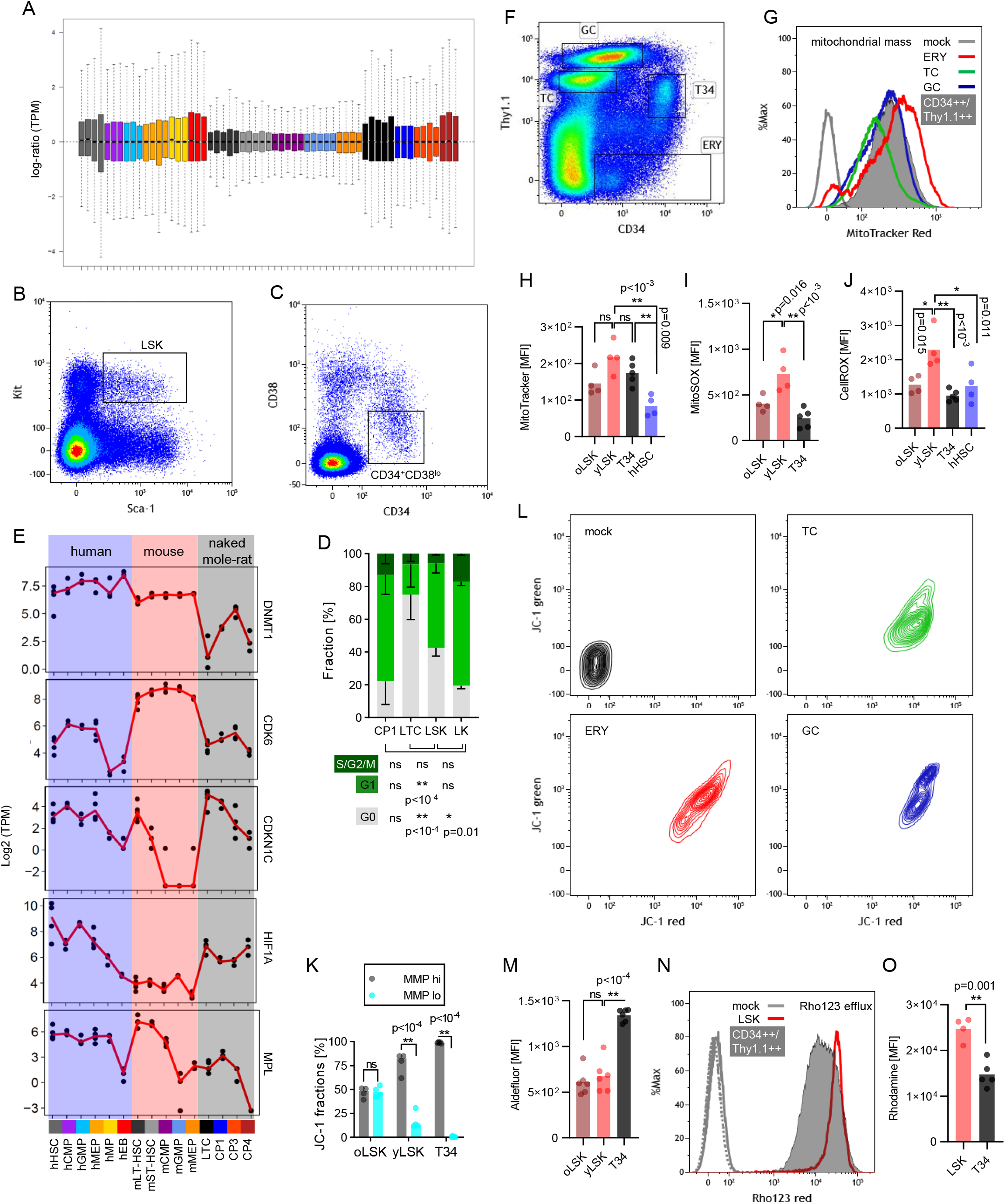

**Figure S7.**
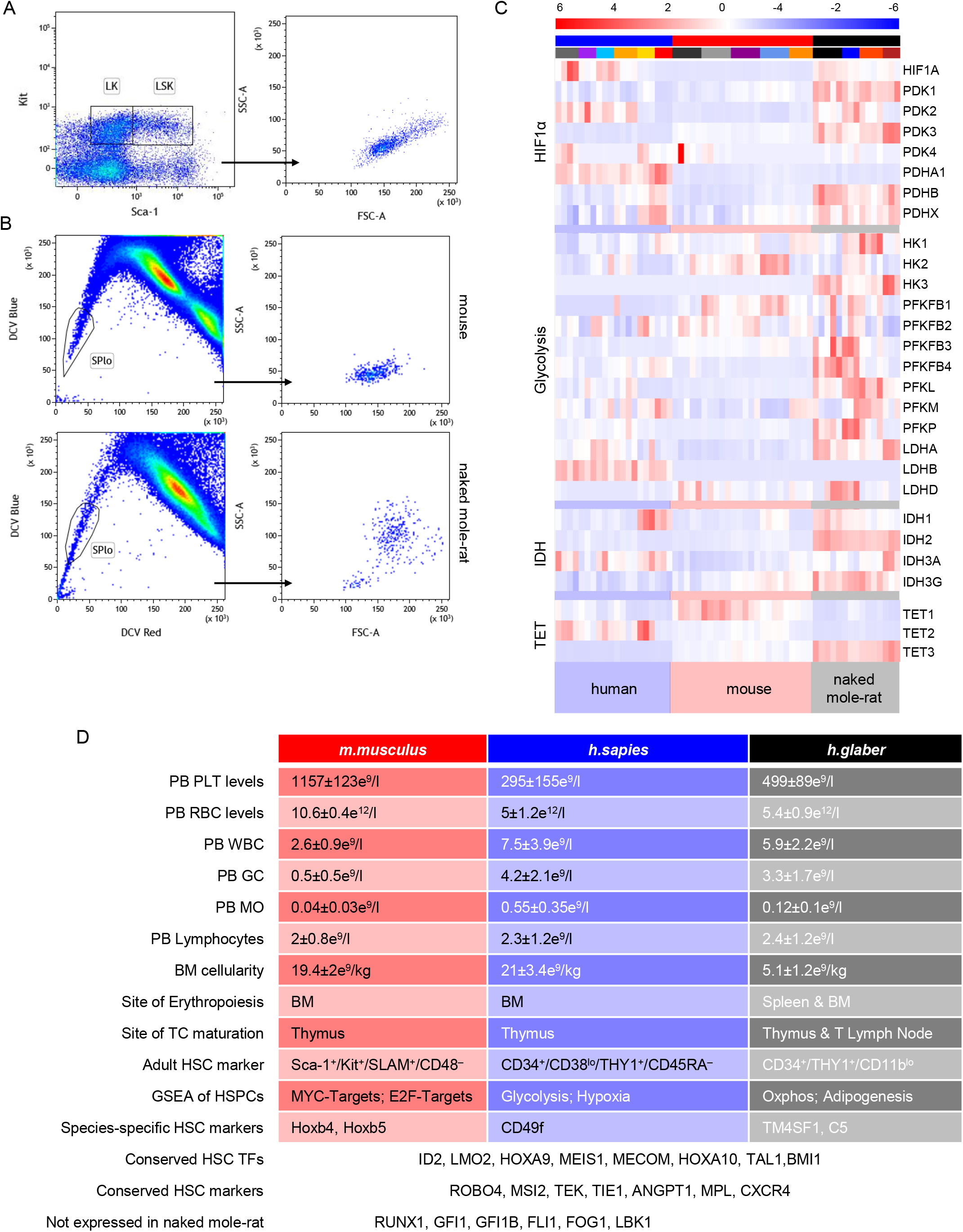

