## Supplementary material for "The hematopoietic landscape at single-cell resolution reveals unexpected stem cell features in naked mole-rats": Methods and Supplemental Figure Legends

### CONTACT FOR REAGENT AND RESOURCE SHARING

### EXPERIMENTAL MODEL AND SUBJECT DETAILS

#### Animals

All animal experiments were approved and performed in accordance with guidelines instructed by the University of Rochester Committee on Animal Resources with protocol numbers 2009-054 (naked mole rat) and 2017-033 (mouse). Naked mole rats were from the University of Rochester colonies, housing conditions as described (Ke et al., 2014). C57BL/6 mice were obtained from NIA, in comparative assays yLSK were sorted from 3-4 month and oLSK from 25 month old mice. Immunodeficient strain NSGS was purchased from JAX.

#### Primary cell isolation

Marrow from mice and naked mole-rats was extracted from femora, tibiae, humeri, iliaci and vertebrae by crushing. Spleen, liver, thymus and lymph nodes were minced over a 70µm strainer and resuspended in FACS buffer. Blood from mice was drawn via retroorbital capillary bleeding, naked mole-rat blood was obtained via heart puncture.

### METHOD DETAILS

#### Hematology Analyzer

PB parameters were measured with a Vet ABC Plus+ (scil) Analyzer. Specifically, naked mole-rat and mouse samples were measured with the “mouse_research” protocol (scil Tech Support, available upon request), which provides a 3-part differential in 17 parameters.

#### Histology

Imaging and analysis was performed using a using a Nikon Eclipse Ti-S microscope. Coverslips were applied with DEPEX Mounting media (Electron Microscopy Sciences), except for Alkaline Phosphatase staining where Vectashield Hard Set Mounting Medium for Fluorescence (Vector) was applied. Femur bones were decalcified with 14% EDTA for a minimum of 2 weeks and stored in 10% neutral buffered formalin. Soft tissues were stored in 10% neutral buffered formalin, processing was done using a Sakura Tissue-Tek VIP 6 automated histoprocessor, paraffin embedding was done using a Sakura Tissue-Tek TEC 5 paraffin embedding center. A Microm HM315 microtome was used to section tissues at a thickness of 5µm, which then were floated onto a slide with a water bath at a temperature between 45°C and 55°C. Sections were deparaffinized and rehydrated to distilled water through xylene and graded ethanol (100% to 70%).

*May-Grünwald-Giemsa*: Cytospins of whole spleen or WBM or sorted cells were prepared using a Rotofix 32A (Hettich) and stained at room temperature with May-Grünwald solution (Sigma) for 5min, washed in phosphate buffer pH 7.2 (Sigma) for 1.5min, and counterstained in 4.8% Modified Giemsa (Sigma) for 13min.

*Alkaline Phosphatase*: Cytospins were stained with the Alkaline Phosphatase kit (Sigma) according to manufacturer's instructions with the exemption of combining FBB-Alkaline Solution with Hematoxylin Solution, Gill No. 3 (Sigma) as counterstain.

*Benzidine Mayer's Hematoxylin*: Slides were fixed at room temperature with methanol for 30sec, incubated with o-Dianisidine (Sigma) 1% in methanol for 1min and stained with $H_2O_2$ 2.5% in ethanol for 30sec before rinsing for 15sec in water and counterstaining with Mayer's Hematoxylin Solution (Sigma) for 2min.

*Wright Giemsa*: Blood films were incubated with Wright-Giemsa Stain (Electron Microscopy Sciences) for 1min, rinsed briefly with water and developed in phosphate buffer pH 7.2 for 2min, then rinsed again. Slides were scored by taking 3 random micrographs of monolayers from the feathered edge of each sample to count both RBCs and platelets and average the technical replicates. Then mean RBC levels from the bloodcounter measurements (mouse, 9.1e$^{12}$/l; naked mole-rat 5.4e$^{12}$/l) were used to convert PLT/RBC ratios to a volumetric PLT count via bloodcounter by $\frac{(RBC\times 10^{12})\times(PLT_{count})}{l\times(RBC_{count})}$

*Hematoxylin & Eosin*: Sections were stained with Mayers Hematoxylin (Sigma) for 1min and washed with tap water to remove excess blue coloring. Soft tissue sections were further decolorized with 3 dips in 0.5% acid alcohol and washed in distilled water. The nuclei of sections were blued in 1X PBS for 1 minute and washed again in distilled water. An Alcoholic-Eosin counterstain was applied for 30sec before slides were immediately dehydrated and cleared through 3 changes of 95% ethanol, 2 changes of 100% ethanol, and three changes of Xylene for 1min each.

*Microwave Giemsa for plastic marrow sections*: Paraffin-embedded Femora were subjected to the microwave modification of a conventional Giemsa stain, which we found to produce clearer contrast of megakaryocytic cells as distinguished by their pale purple cytoplasm and abundant nuclear chromatin staining due to polyploidy. The stain was performed as described in www.urmc.rochester.edu/urmc-labs/pathology. Slides were scored by taking 3 random micrographs of marrow from medullary canal for each sample to count polyploidy giant Megakaryocytes, average the technical replicates and convert micrograph pixel size via magnification to bone area in $mm^2$.

#### Global methylation levels

Trizol-extracted DNA from sorted mouse and naked mole-rat BM cells were used for the MethylFlash ELISA (Epigentek). 50ng DNA per sample replicate was used as input, a standard curve prepared from the kit components was used to calculate % of methylation by linear regression as decribed in the manufactures instructions.

#### Total BM cellularity

Human total BM cellularity was taken from (Skarberg, 1974). Mouse total BM cellularity was taken from (Colvin et al., 2004) using the data from an 8 month C57BL/6J and the corresponding body weight from the JAX lab (Stock No. 000664). Total live cell counts from naked mole-rat BM extractions were obtained by Trypan-Blue exclusion dye counts on a Neubauer improved Hemcytometer (Marienfeld). According to (Colvin et al., 2004) the marrow spaces of femora, tibiae, humeri, iliaci and vertebrae comprise 78% of total marrow in a 2 month BALB/c mouse. As there is no comparable information for naked mole-rats available, we used this figure as an estimate to scale the mean BM cellularity $129.6e^6 \pm 38.8$ of 28 animals 10-18 months of age, then using mean body mass 32.6±5.7g of the same cohort to calculate total BM cellularity.

### Methylcellulose colony assays

Fresh sorted or whole BM naked mole-rat cells were tested to grow in mouse (M3434, SCT), rat (R3774, SCT) or human (H4435, SCT or HSC005, RnD Systems) methylcellulose formulations to show the highest colony numbers, colony sizes and cell viability with human cytokine cocktails. Unless otherwise stated, $2x10^3$ sorted naked mole-rat cells were added to 3ml of HSC005 supplemented with 1% Penicillin/Streptomycin and 1X GlutaMAX (both Thermo Fisher), grown for 14d at 32°C, 5% $CO_2$ and 3% $O_2$, and scored. Although hematopoietic naked mole-rat cells will grow at 37°C, the total number as well as colony and cell type diversity is strongly enhanced at 32°C (data not shown). Colony assays grown at 37°C give rise to two types of colonies (erythroid vs myeloid), which are notably smaller than each of the 4 colony types we can distinguish at 32°C (Figure S3J). Replatings were done by resuspending scored dishes at day 14 in FACS buffer, count cells and replate $1x10^4$ cells into above growth conditions. CP3 cells did not grow substantially in the first replating, and a second replating had no sizable colonies for all naked mole-rat HSPC types. Benzidine staining of naked mole-rat methylcellulose assays was done as described (Murphy, 1978). Briefly, a 0.2% benzidine dihydrochloride (Sigma) solution in 0.5M acetic acid was prepared, which was supplemented with 0.24% of 50% $H_2O_2$. 1ml of this solution was layered carefully over each dish, and past 5min colonies were scored for the proportion of colonies which are uniformly benzidine-unreactive (color-less), uniformly benzidine-reactive (blue) and colonies containing both reactive and unreactive cells (mixed colonies containing both differentiated, hemoglobin containing and non-erythroid cells).

### Flow Cytometry

Flow cytometry analysis was performed at the URMC Flow Core on a LSR II or LSRFortessa (both BD). Kaluza 2.1 (Beckman Coulter) was used for data analysis. Staining and measurement were done using standard protocols. Red blood cell lysis was done by resuspending BM pellets in 4ml, spleen pellets in 1ml and up to 500µl PB in 20ml of RBC lysis buffer, prepared by dissolving 4.1g $NH_4Cl$ and 0.5g $KHCO_3^-$ into 500ml double-distilled $H_2O$ and adding 200µl 0.5M EDTA. BM and spleens were incubated for 2min on ice, PB were lysed for 30min at room temperature. Cells were resuspended in FACS buffer (DPBS, 2mM EDTA, 2% FBS [Gibco]) at $1x10^7$ cells/ml, antibodies were added at $1µl/10^7$ cells, vortex-mixed and incubated for 30min at 4°C in the dark. The primary gating path for all unfixed samples was: scatter-gated WBC (FSC-A vs SSC-A) => singlets1 (SSC-W vs SSC-H) => singlets2 (FSC-W vs FSC-H) => viable cells (SSC vs DAPI) == proceed with specific markers/probes. Immunophenotyping of naked mole-rat BM, spleen, thymus, PB and lymph nodes: CD90 FITC; CD125 PE; Thy1.1 PE-Cy7; CD34 APC, CD11b APC-Cy7. Quantification of murine BM SLAM HSCs was performed using mouse LIN Pacific Blue; Sca-1 BUV395; CD150 PE; Kit PE-Cy7; CD48 APC-Cy7. Quantification of human BM LT-HSCs was performed using human LIN Pacific Blue; CD34 APC; CD38 APC-Cy7; CD45RA FITC; CD90 PE-Cy7.

*Sorting* was performed at the URMC Flow Core on a FACSAria (BD) using a 85µm nozzle, staining was done as described. Human HSCs were sorted for population RNA-Seq as $LIN^-$/$CD34^+$/$CD38^{Lo}$/$CD45RA^-$/$CD90^{Dim}$. Naked mole-rat HSPC populations were sorted as described with a lineage cocktail comprised of CD11b, CD18, CD90 and CD125 (NMR LIN). Naked mole-rat BM and spleen sorting panel was: NMR LIN Pacific Blue; Thy1.1 PE-Cy7; CD34 APC. Naked mole-rat PB sorting panel was: Thy1.1 PE-Cy7; CD11b APC-Cy7.

*Molecular probing* was performed on frozen aliquots from mouse and naked mole-rat BM. For each probe, cells were diluted in 1ml pre-warmed DMEM+ at $1x10^6$ cells/ml. All stainings were performed simultaneously for 4-6 naked mole-rat, 2-4 old, 2-4 young mice and 4 human biological replicates. ALDEFLUOR (SCT) reagent was added at 0.5µl/ml, mixed and incubated for 15min at 37°C in a water bath. MitoTracker Orange CMTMRos (Thermo Fisher) was added at 10nM and incubated for 45min at 37°C; MitoSOX red (Thermo Fisher) was added at 5µM and incubated for 30min at 37°C; MitoStatus TMRE (BD) was added to $0.5x10^6$ cells/ml with 25nM and incubated for 10min at room temperature in the dark; JC-1 (Thermo Fisher) was added at 1µM with or without 5µM FCCP and incubated for 15min at 37°C; CellROX Orange (Thermo Fisher) was added at 5µM and incubated for 60min at 37°C. The subsequent antibody staining was performed as above with 30min incubation on ice, panel was Sca-1 (mouse) or CD34 (naked mole-rat) APC; Kit (mouse) or Thy1.1 (naked mole-rat) PE-Cy7; Lineage Cocktail V450; 1µg/ml DAPI (Thermo Fisher) was used as viability stain.

*Pyronin Y* staining was performed on frozen aliquots from mouse and naked mole-rat BM. Briefly, cells were count-adjusted to $1x10^6$ cells/ml and resuspended into 1ml of DMEM+ (DMEM high Glucose, 2% FBS, 10mM HEPES; all Gibco). Upon addition of 50µg/ml Verapamil (Sigma) and 5µM DyeCycle Violet (Thermo Fisher) cells were incubated for 45min at 37°C in a water bath, vortex-mixed every 15min. Past 45min 0.1µg/ml Pyronin Y was added to the reaction and incubated an additional 15min at 37°C, then washed with 3ml ice-cold Staining buffer (HBSS [Gibco], 0.33M HEPES, 3.5% FBS, 0.02% $NaN_3$ [Sigma]). A subsequent antibody staining was performed as above with incubation on ice, panel was Sca-1 (mouse) or CD34 (naked mole-rat) APC; Kit (mouse) or Thy1.1 (naked mole-rat) APC-Cy7; Lineage Cocktail FITC; 500nM SYTOX green was used as viability stain.

*Ki67* staining was performed on frozen aliquots from mouse and naked mole-rat BM. Cells were count-adjusted to $1x10^7$ cells/ml and antibody staining was performed as described, panel was Sca-1 (mouse) or CD34 (naked mole-rat) APC; Kit (mouse) or Thy1.1 (naked mole-rat) APC-Cy7; Lineage Cocktail FITC. For fixation and permeabilization we used the buffers from the BrdU Flow Kit (BD). Briefly, cells were fixed for 30min in Cytofix/Cytoperm on ice at $100µl/1x10^6$ cells, permeabilized for 10min in CytopermPlus on ice at $100µl/1x10^6$ cells, refixed for 5min in Cytofix/Cytoperm on ice at $100µl/1x10^6$ cells, all washes done with 1X Perm/Wash. Cells were resuspended in Staining buffer at $1x10^7$ cells/ml, Ki67 antibodies (mouse: clone 16A8; naked mole-rat: clone Ki-67; both PE-conjugated, BioLegend) were added at $5µl/1x10^6$ cells and incubated for 30min at room temperature in the dark, 1µg/ml DAPI was used as DNA stain.

*Rhodamine 123* staining was performed by incubating 1x$10^6$ cells for 30min with 1µg/ml Rho in HBSS+ (HBSS, 2% FBS, 10mM HEPES; all Gibco) at 37°C, then cells were washed with 2ml HBSS+, spun down and reincubated for 15min at 37°C, 1µg/ml DAPI was used as DNA stain. *Side population (SP)* staining was done as described previously (Goodell et al., 1996; Telford et al., 2007) with using DyeCycle Violet (DCV) instead of Hoechst 33342 on frozen aliquots from mouse and naked mole-rat BM. Cells were resuspended in pre-warmed DMEM+ at 1x$10^6$ cells/ml and incubated with 5µM DCV in 1ml for 15, 30, 45, 60, 75, 90, 105 or 120min. All washes and subsequent antibody staining were performed on ice in HBSS+ (HBSS, 2% FBS, 10mM HEPES; all Gibco), panel was Sca-1 (mouse), CD34 (naked mole-rat) APC; Kit (mouse), Thy1.1 (naked mole-rat) APC-Cy7; Lineage Cocktail FITC, 0.5µg/ml PI (Roche) was used as viability stain. While we detected earliest emergence of mouse SP at 45min, SP frequency peeks between 90 and 105min (data not shown). By contrast, naked mole-rat BM cells show no SP at 75min and later time points, while 45 and 60min gave rise to >10-fold less cells in the SP area than in murine cells (Figure S7, data not shown). Moreover, we run the same time course on rat and guinea pig BM cells to find a similar pattern than for naked mole-rats, with an infrequent and inconsistent SP which is distributed over all three major blood cell types distinguishable by FSC-SSC backgating (data not shown). We verified the emergence of mouse SP by adding 50µM Verapamil alongside DCV, which strongly impaired SP formation at 90 and 120min of incubation (data not shown).

**Xenotransplantations**

Naked mole-rat BM and/or spleen cells were extracted, sorted and directly transplanted into 2.5Gy-irradiated (24h pre Tx) NSGS recipients between 5-9 weeks of age at cell doses between 50-200k cells. Injections were done via the retroorbital sinus, blood sampling was performed via maxillary vein or retroorbital plexus at weeks 4, 8 and 12. Hosts were culled at 2, 4, 8 or 12 weeks and engraftment frequencies were estimated by flow cytometry using only naked mole-rat markers not cross-reactive with mouse cells and CD45.1 (A20, BioLegend), then engraftment rates were adjusted for input cell dose to 100k/Tx. Gating path was WBC (FSC-A vs SSC-A) => singlets1 (SSC-W vs SSC-H) => singlets2 (FSC-W vs FSC-H) => viable cells (SSC vs DAPI) => NOT Thy1.1$^{-}$/CD34$^{-}$ (CD34 vs Thy1.1) == engrafted naked mole-rat cells (Figure 4G). One limitation for quantifying engraftment levels is that naked mole-rat BM features cells negative for the above markers which can arise from transplanted HSPCs as xenogenic CP7 (Figure 3D, S3H). A cross-reactive guinea pig CD45 antibody does not stain >80% of naked mole-rat WBM cells and exhibits notable cross-reactivity with BM from NSGS recipients (Figure S3I). We further detected cells double-positive for guinea pig CD45 and CD45.1 (Figure S3L). Cells stained as Thy1.1$^{+}$ and/or CD34$^{+}$ are clearly originated by the xenograft as untransplanted NSGS BM does not feature any Thy1.1 of CD34 labelled cells (Figure S3H). Since all three different cell populations from guinea pig CD45 vs CD45.1 staining (DN, CD45.1+, CD45+/CD45.1-) contain a different pattern of cells stained as Thy1.1$^{+}$ and/or CD34$^{+}$, we considered any cell positive for one or both markers as xenograft. We reasoned that due to the in vitro cross-reactivity of human SCF engraftment would be supported when using NSGS hosts. However, when we compared the engraftment efficiency for ~1x10$^{5}$ LTCs transplanted into NSGB (NOD.Cg-*B2m*$^{tm1Unc}$ *Prkdc*$^{scid}$ *Il2rg*$^{tm1Wjl}$/SzJ) or NSGS mice at 4 weeks and same cell dose between NSG (NOD.Cg-*Prkdc*$^{scid}$ *Il2rg*$^{tm1Wjl}$/SzJ) and NSGS at 8 weeks, we found no significant differences between the strains (data not shown).

**Transcriptome assembly**

All Naked mole-rat RNA-Seq was performed with the GRC URMC Rochester. RNA from WBM was sequenced with ~230 million reads on a HiSeq2500v4 (Illumina). Raw Illumina paired-end sequencing reads where assessed with FastQC. Rcorrector (Song and Florea, 2015) was used to correct sequencing errors and read pairs with uncorrectable errors were removed using a custom python script (GRC URMC Rochester). Adapter and base quality trimming was performed using trim galore and cutadapt (Martin, 2011) resulting in high quality reads that were used as input to Trinity (Grabherr et al., 2011) for assembly. FRAMA (Bens et al., 2016) was used to post-process the *de novo* assembly, including reduction of contig redundancy, ortholog assignment using human as a reference, correction of misassembled transcripts, scaffolding of fragmented transcripts and coding sequence identification. Quality assessment of the final FRAMA transcriptome was performed using BUSCO and TransRate (Simao et al., 2015; Smith-Unna et al., 2016). The transcriptome was mapped by blastn to the naked mole-rat genome (hetgla_female_1.0) or to transcript sequences annotated in ENSEMBL97. Mapped genomic coordinates of transcripts were thus compared to those of annotated genes using a custom python script. We found that 512 non-overlapping FRAMA transcripts (i.e., gene loci) were absent from the annotation, and another 5281 had >20% transcript length mapped to the genome but not matching annotated isoforms (Figure S1I-J, Table S1). Interestingly, most of the novel genes do not overlap with genes from hematopoietic stem and progenitor cell genesets (Figure S1K) (Schwarzer et al., 2017).

**Population RNA-Seq**

RNA from sorted human and naked mole-rat populations was sequenced at ~100 million reads on a HiSeq2500v4 (Illumina), the ClonTech v4 kit (Takara) was used for library preparation. Raw Illumina paired-end sequencing reads where assessed with FastQC. Adapter and base quality trimming was performed using Trimmomatic (Bolger et al., 2014) resulting in high quality reads that were assessed again with FastQC. *RSEM v1.3.0* was used to calculate expected counts and TPMs (Li and Dewey, 2011). We used a customized perl script to run RSEM with the FRAMA transcriptome as reference using *bowtie2* aligner option. We also run RSEM with the ENSEMBL94 hetgla_female_1.0 annotation using *STAR* aligner option to confirm all clusterings and differential gene expression signatures for all naked mole-rat samples, results were almost identical to data shown in Figures 2, 3, 6 and 7 (data not shown).

Subsequent analysis was done with *R 3.6.1*and *Bioconductor* (Gentleman et al., 2004); *edgeR* was used to calculate size factors with `method="RLE"` and computing CPMs (Robinson et al., 2010). We applied *genefilter* to calculate the interquartile range (IQR) of CPMs with `IQR(x) > 1` to filter unexpressed and outlier genes; libray-size normalized, IQR-filtered log2-transformed CPMs were shown in Figure S2I, S1L, S1Q, S6H. CPMs were vst-transformed by *DESeq2* (Love et al., 2014), then a PCA from stats package was used as input for Rtsne (van der Maaten and Hinton, 2008). We applied *limma* to perform voom-transformation and select for differentially expressed genes (DEGs) with $p < 0.05$ and log-fold-change 1 (Ritchie et al., 2015). GSEA was performed using the *gsva* package with `method="ssGSEA"` using either the hematopoietic stem and progenitor geneset collection as described in (Schwarzer et al., 2017) or the MSigDB v6.0 hallmark genesets with a p-value threshold of 0.05 (Barbie et al., 2009; Hanzelmann et al., 2013; Liberzon et al., 2015; Subramanian et al., 2005); all GSEA calculations were performed on the combined up- and downregulated DEG signature for each group. The *fGSEA* package was used to retrieve leading edge genes after reperforming GSEA under default conditions (Sergushichev, 2016), required gene rank metric was generated after (Plaisier et al., 2010).

For Figure 3A and following we trimmed the dataset from 19 to 15 samples based on the least common multiple to retain highest replicate number per group to form a separated cluster in the t-SNE.

The expression gradient was calculated by a customized R function, which ordered the log2-transformed CPMs for each gene along their numeric value, allowing to filter out the genes subsequently changing expression from one group to another.

We validated our preprocessing, clustering and differential gene expression (DGE) pipeline by using previously reported human and mouse RNA-Seq datasets of hematopoietic stem and progenitor cells. The human dataset from (Chen et al., 2014) was accessed via the BLUEPRINT consortium under EGAD00001000745 and included transcriptomes of HSCs, common myeloid progenitors (CMPs), granulocytic/monocytic progenitors (GMPs), megakaryocytic/erythroid progenitors (MEPs), megakaryocytic precursors (MPs) and erythroblasts (EBs) (Figure S2N). The pulled data consisted of RSEM counts and TPMs generated by the BLUEPRINT RNA-Seq analysis pipeline. The cleaned dataset comprised 14080 genes with HGNC identifiers (Braschi et al., 2019) detected across 20 hematopoietic samples (Figure S2O). T-SNE clustering faithfully divided progenitors from more mature precursors and blasts (Figure S2P). Moreover, HSCs were clearly separated from other samples, likewise GMPs formed an individual group, while there was residual overlap between MEP and CMP transcriptomes, most probably reflecting the transcriptional plasticity of CMPs (Novershtern et al., 2011). The number of DEGs being lowest in CMPs and highest in the more distant precursors also points towards a continuum of cell states with CMPs as the hub between stem cells and more directed progenitors (Figure S2Q). We then run single sample gene set enrichment analysis (ssGSEA) against a collection of 81 HSPC signatures (Schwarzer et al., 2017). We chose 5 genesets of any cell type with the lowest q-value to consolidate the developmental annotations (Figure S2R). Strikingly, this precisely matched the *a priori* cell type with its corresponding genesets for HSCs, MPs and EBs. Although most displayed genesets are derived from human datasets, there are erythroid, myeloid, and lymphomyeloid signatures from murine datasets among the top 5 genesets, demonstrating the capacity for cross-species annotation of hematopoietic cell types (Figure S2R).

The murine dataset created by the Jackson Laboratory (George et al., 2016) was pulled, trimmed, aligned and read-counted from GEO GSE74691 by a custom pipeline using *sratoolkit, cutadapt, trimgalore, trimmomatic* and *RSEM* with *STAR* aligner against ENSEMBL94 GRCm38. The dataset comprised LT- and ST-HSCs, CMPs, GMPs and MEPs (Figure S2S). The processed dataset had 12477 HGNC genes expressed over 24 samples, clustering separated all progenitor groups from each other and HSCs, however since LT- and ST-HSCs overlapped we combined them to a joint HSC cluster (Figure S2T-V). GSEA recapitulated the transcriptional annotation of the human dataset for HSCs (Figure S2W). No significantly enriched geneset could be obtained for CMPs due to only 16 DEGs for this group, supporting the CMPs central position in the hematopoietic hierarchy (Figure S2V-W).

The 3 species comparison utilized the BLUEPRINT dataset from Figure S2K-O, the JAX GEO dataset from Figure S2P-T and the naked mole-rat dataset from Figure S2I. Human, mouse and naked mole-rat datasets (Figure 7) were merged based on HGNC symbols, then *genefilter* was used to calculate IQR of TPMs with `IQR(x) > 1` to filter unexpressed and outlier genes; The *TCC* package was used to calculate TMM-based size-factors (Sun et al., 2013). The function `betweenLaneNormalization` with median scaling from the *EDASeq* package was used to normalize for sequencing batch effects (Risso et al., 2011); libray-size and batch normalized, IQR-filtered log2-transformed TPMs were shown in Figure S6K-L. TPMs were vst-transformed by *DESeq2*, then a PCA from stats package was used as input for Rtsne; DGE and GSEA were performed as above.

**Single cell RNA-Seq**

BM, blood and thymus cells from a male and a female animal aged 11 months were enriched by sorting. For BM we sorted CP1 3k, LTC 3k, CP3 2k, CP4 2k, CP5 2k, CP6 2k, CP7 3k; total 30000 marrow cells from 2 animals. For PB we sorted GC 1k, MO 1k, BC 1k, TC 1.5k, EO 0.5k; total 10000 peripheral blood leukocytes from 2 animals. For thymus we sorted ETP 1k, TSP 1k, LTC 1k; total 6000 thymocytes from 2 animals. Cells were pooled according to their tissue origins and processed for CITE-Seq using a protocol from the Stoeckius lab and the Chromium Single-Cell 3′ Library (Stoeckius et al., 2017). Raw reads generated on the Illumina NovaSeq6000 sequencer were demultiplexed using *Cell Ranger 3.0.2* software in conjunction with Illumina's *bcl2fastq 2.19.0*. Cell Ranger was also used to align the read data to the FRAMA *de novo* transcriptome assembly and ENSEMBL94 hetgla_female_1.0, barcode count, UMI compress, and filter for "`true`" cells. CITE-Seq data for each capture was also demultiplexed using bcl2fastq and processed with CITE-seq-Count 1.4.2 (Roelli et al., 2019) given the antibody barcode sequences, a white list of filtered cell barcodes from the matching Cell Ranger "`count`" run, and parameters: "`-cbf 1 -cbl 16 -umif 17 -umil 26`".

Subsequent analysis was done with *R 3.6.1* and *Bioconductor*. 10X files were assigned to a *SingleCellExperiment* S4 class *(Lun A, 2019)*, and each gene without any counts in any cell was immediately removed. We removed every cell with > 99% 0's, and every gene not expressed in > 10 cells, according to (Morgan Mike and Ballereau Stephane, 2018). The *scran* package was applied to compute size-factors with "`min.mean=0.1`" (Lun et al., 2016), finally *scater* normalized the *SingleCellExperiment* object (McCarthy et al., 2017).

We converted the S4 class into a *Seurat* object and added the CITE-signals as described in the satija-lab tutorial (Butler et al., 2018; Stuart et al., 2019). The *Seurat* object contained the transcriptomic data and the CITE-Seq counts each in form of an independent "`assay`"; RNA assay was log-normalized with "`scale.factor = 1e4`", CITE assay was "`CLR`" normalized.

Variable features were detected with arguments `selection.method = "vst", nfeatures = 2000, mean.cutoff = c(0.1, 10); nfeatures = 3000` for the combined dataset (Figure 5).

We then performed a cell cycle scoring as described in the respective *Seurat* vignette. For PB (Figure 1F) and thymus (Figure 4M) libraries PCA was run on the unregressed objects variable features. For BM (Figure 4A) library we performed a regression of the difference between G2M and S phase scores, leaving signals separating non-cycling and cycling cells intact (see vignette). For the combined dataset (Figure 5A) we applied the complete cell cycle regression model.

The data was scaled using *Seurat's* `ScaleData` function with `center = TRUE, scale = TRUE`.

The DEGs for each cluster were detected by `FindAllMarkers` function with arguments `test.use = "MAST", logfc.threshold = log(2), min.pct = 0.25, return.thresh = 0.05`.

Clustering was done using *Seurat's* `FindClusters` function with `resolution = 0.5` for PB, BM and the combined dataset; THY was done with `resolution = 0.1`. Higher resolution clustering yielded more thymic clusters, however positive (upregulated) markers were not obtained for all clusters and mostly reflected cell cycle state and housekeeping genes, whereas upon cell cycle regression clustering was less effective and uninformative (data not shown). Given the low complexity of droplet-based libraries (1758 median genes/cell for thymus), scRNA-Seq with higher coverage for future follow-ups is warranted.

The BM clustering featured 16 clusters, of which 15 could be annotated to cell types based on *fGSEA* or curated marker expression (Figure 4A-F, S4F-I, Table S2-3). We also detected a cluster mp (multiplets) with 5 marker genes (S100A9, S100A10, S100A4, LSP1, AP1S2), which are also found in $CD14^+$ monocytes, $CD8a^-$ DCs, $CD34^+$ cells, $CD33^+$ myeloid cells, Tregs and macrophages. The mp cells co-cluster with ERY5 in UMAP dimensionality reduction, and their signature is not retained in the mp cluster of the combined dataset (Table M1).

**Table M1: Total cell numbers per cluster for each dataset alone or combined**

| PB | | THY | | BM | | COMBINED | | COMBINED.no.mp | |
|---|---|---|---|---|---|---|---|---|---|
| *cluster* | cells | *cluster* | cells | *cluster* | cells | *cluster* | cells | *cluster* | cells |
| | | | | HSPC1 | 2635 | HSC | 3533 | HSC | 3533 |
| | | | | HSPC2 | 1440 | MPP | 1503 | MPP | 1413 |
| | | | | HSPC3 | 871 | BCP | 552 | BCP | 714 |
| BC1 | 555 | | | BC1 | 533 | BC | 978 | BC | 978 |
| BC2 | 12 | | | BC2 | 46 | | | | |
| | | | | BC3 | 422 | | | | |
| | | | | PC | 47 | BM.PC | 50 | BM.PC | 50 |
| | | TSP | 933 | | | TSP | 1563 | TSP | 1558 |
| | | TC | 685 | | | | | | |
| | | ETP | 848 | | | ETP | 833 | ETP | 838 |
| CD4-TC | 617 | | | | | PB.TC | 910 | PB.TC | 913 |
| CD8-TC | 37 | | | | | | | | |
| TC | 180 | | | | | | | | |

| | | | | | | | |
|---|---|---|---|---|---|---|---|
| | | MY | 386 | BM.MY | 369 | BM.MY | 270 |
| DC | 57 | | | PB.DC | 60 | PB.DC | 60 |
| MO | 608 | | | PB.MO | 636 | PB.MO | 636 |
| GC | 528 | | | PB.GC | 687 | PB.GC | 686 |
| | | HSPC4 | 637 | MEP | 649 | MEP | 656 |
| | | ERY1 | 1722 | ERY1 | 2787 | ERY1 | 2899 |
| | | ERY2 | 1203 | | | | |
| | | ERY3 | 1515 | ERY2 | 1641 | ERY2 | 1526 |
| | | ERY4 | 955 | ERY3 | 959 | ERY3 | 959 |
| | | ERY5 | 346 | ERY4 | 321 | ERY4 | 321 |
| | | SC | 20 | BM.SC | 23 | BM.SC | 23 |
| | | | | | | ua | 21 |
| | | mp | 136 | mp | 78 | | |

The clustering of the combined dataset retrieved 19 clusters, of which 18 could be annotated to hematopoietic cell types based on *fGSEA* or retention of a cluster signature from the PB, thymus or BM datasets (Table S6). A cluster mp with 220 specifically upregulated genes did not feature any markers from the BM dataset mp cluster, and UMAP located mp in the vicinity to BCPs and PB.TCs. This mp cluster exclusively yielded terms with NES < 0 upon *fGSEA* with the hematopoietic geneset collection (see below), while an input of the 220 upregulated genes into the *Enrichr* tool (Kuleshov et al., 2016) found the top two significant cell type associations from the human gene atlas to be B-Lymphoblasts and CD34+ cells ($q<2.9e^{-9}$; $q<8.3e^{-3}$), top two associations from mouse gene atlas were MEP and embryonic stem cell line Bruce4 ($q<2.8e^{-5}$; $q<1.9e^{-3}$), top two associations from ARCHS4 tissue database were Kidney bulk and CD34+ cells ($q<3e^{-19}$; $q<1.1e^{-2}$). For pseudotime and trajectory computations we removed the mp cluster from the combined dataset to avoid potential interference of functionally unassociated cells, then re-clustered with same parameters (Table M1). Post mp cluster removal re-clustering essentially restored all major clusters with slight differences in cell numbers per cluster (Table M1). The ua (unassigned) cluster with 21 cells featured 4 significantly upregulated markers, all of which are also HSC markers (Table S4).

Hematopoietic cell type annotation was done based on single markers conserved for broad cell types, such as CD4 to CD4-TCs, or *fGSEA* was applied using the HSPC geneset collection from (Schwarzer et al., 2017) with the upregulated DEGs from the population RNA-Seq analysis (Table S5 sheet 'hema.GENESETS.hsa.mmu.hgl') added from this study. To determine the rank metrics for *fGSEA* the q-value requires to be transformed by `-log10(q-value)`; (Plaisier et al., 2010). *Seurat's* `FindAllMarkers` function can generate 0 q-values (p_val_adj, Table S4) for high confidence hits, thus for any 0 we added the lowest q-value > 0 of the entire marker list for the group to test to each marker with q-value = 0. For the BM partitions (Table S3) we chose all terms with NES > 0 and padj < 0.05 (q-value), except for the MY group with markers < 150, where we filtered by NES > 0 and pval < 0.05 (statistical significance of GSEA tools correlates with number of input genes matched to terms); The SC cluster did not retrieve any HSPC cell type term with *fGSEA* and was not included in any of the 4 partitions. For the clusters from the combined dataset (Table S5) we chose all terms with NES > 0 and padj < 0.05; The clusters BM.MY, BM.PC had no significant terms, clusters MEP and mp only terms with NES < 0.

The signature of BM HSPC1 vs HSPC2 (Table S4) was fed into the *Enrichr* webtool (Kuleshov et al., 2016), Table S5 sheet 'MPP.vs.HSC.UP.mitotic.genes' shows the top 20 Combined Score pathways from the *Enrichr* Reactome 2016 pathway categories.

For pseudotime and trajectory computations the *dynverse* package (Saelens et al., 2019) was applied with inputs of the Louvain cluster id's for each cell along with one `start_id` (HSCs) and 5 `end_id` (BC, PB.TC, PB.MO, PB.BC, ERY4) states. We used an adaptation of the PAGA algorithm (Wolf et al., 2019) for `infer_trajectory` with `method = ti_Paga_tree`, `embedding_type = "umap"` and defaults. For the dendrogram a random HSC cell id was specified as root. Alternatively, for MST-based trajectory we used `infer_trajectory` with `method = ti_mst`, and specified the same root as with the PAGA computation for the dendrogram.

The *ouija* package (Campbell and Yau, 2019) was used to compute a descriptive single cell pseudotime using logcounts and `inference_type = "vb"`. We designated markers of committed progenitors and mature cells as switch genes, markers of the primitive cell types HSC, MPP and BCP were set as transient. We then assigned a prior switch strength of -10 to all erythroid cell fate-associated markers: `c("EPOR", "TPM1", "TSPO2", "TFRC", "STOM", "RHAG", "RHD", "MLLT3", "FECH", "GATA1", "HEMGN", "HBB")`. Accordingly, a switch strength of 10 was applied to all myeloid and mature TC and SC markers: `c("CXCR4", "CD4", "CD3E", "CD8B", "ZEB1", "TLR7", "SPARC", "BGLAP", "SIGLEC5", "SELL", "MS4A1", "LYZ", "CD44", "JUNB", "SPI1", "CEBPA")`. A prior `switch_strength_sd = 0.5` was set for all switches. The transient genes were chosen based on known roles in proliferation, HSC to MPP/LMPP transition or receptors and TFs from MAST analysis: `c("VIM", "TOP2A", "TGFB1", "TM6SF1", "SPINT2", "SOX4", "SEPP1", "PYCARD", "MYC", "MSI2", "MDK", "LY6E", "HOXA9", "CD34", "IL2RG", "KLF6", "TM4SF1", "EPCAM", "HHEX", "FOS", "GATA3" ,"TCF7", "ID2")`. These presettings allowed ordering of the multifurcated early hematopoietic hierarchy into a central primitive HSCP block flanked by either erythroid or lymphomyeloid lineages (Figure 5F). In this model, ouija metastable cell states do not reflect a priori cell types but rather outline contingents of each clusters transcriptome to a linear projection of cell states (Figure S6A). Thus almost no erythroid transcriptomes are present in the first three states, accordingly there is strongly diminished transcriptome association with the last four states. HSCs, MPPs and BCPs are broadly spread over states 3-7, with a inverted HSC/MPP ratio present in states 2 vs 8, probably due to the fact that HSCs directly convert to MEPs, while MPPs are the immediate transition into BCPs (most likely the naked mole-rat LMPP fraction) and myeloid progenitors (Figure S6).

The *destiny* package (Angerer et al., 2016) was applied to run diffusion maps with defaults using the logcounts generated by the *SingleCellExperiment* object during the data cleaning, the first eigenvector was then plotted as pseudotime (Figure 5L).

### QUANTIFICATION AND STATISTICAL ANALYSIS

Data are presented as the mean ± SD. Statistical tests performed can be found in the figure legends. P values of less than 0.05 were considered statistically significant. Statistical analyses were carried out using Prism 8 software (GraphPad) unless otherwise stated.

STEMCELL Technologies. (DOCUMENT #28048 | VERSION 3.0.0). Frequencies and Percentages of Mouse Immune Cell Types.

STEMCELL Technologies. (DOCUMENT #23629 | VERSION 4.0.0). Frequencies of Cell Types in Human Peripheral Blood.

Stoeckius, M., Hafemeister, C., Stephenson, W., Houck-Loomis, B., Chattopadhyay, P.K., Swerdlow, H., Satija, R., and Smibert, P. (2017). Simultaneous epitope and transcriptome measurement in single cells. Nat. Methods *14,* 865-868.

Stuart, T., Butler, A., Hoffman, P., Hafemeister, C., Papalexi, E., Mauck, W.M., Hao, Y., Stoeckius, M., Smibert, P., and Satija, R. (2019). Comprehensive Integration of Single-Cell Data. Cell *177,* 188-1902.e21.

Subramanian, A., Tamayo, P., Mootha, V.K., Mukherjee, S., Ebert, B.L., Gillette, M.A., Paulovich, A., Pomeroy, S.L., Golub, T.R., Lander, E.S., and Mesirov, J.P. (2005). Gene set enrichment analysis: a knowledge-based approach for interpreting genome-wide expression profiles. Proc. Natl. Acad. Sci. U. S. A. *102,* 15545-15550.

Sun, J., Nishiyama, T., Shimizu, K., and Kadota, K. (2013). TCC: an R package for comparing tag count data with robust normalization strategies. BMC Bioinformatics *14,* 21-219.

Telford, W.G., Bradford, J., Godfrey, W., Robey, R.W., and Bates, S.E. (2007). Side population analysis using a violet-excited cell-permeable DNA binding dye. Stem Cells *25,* 1029-1036.

van der Maaten, L., and Hinton, G. (2008). Visualizing Data using t-SNE. J. Mach. Learn. Res. *9(Nov),* 2579-2605.

Wolf, F.A., Hamey, F.K., Plass, M., Solana, J., Dahlin, J.S., Gottgens, B., Rajewsky, N., Simon, L., and Theis, F.J. (2019). PAGA: graph abstraction reconciles clustering with trajectory inference through a topology preserving map of single cells. Genome Biol. *20,* 5-x.

### SUPPLEMENTAL FIGURE LEGENDS

**Figure S1. FACS antibody staining pattern validation, transcriptome assembly statistics and blood CITE-Seq QC, Related to Figure 1**

Successive gating scheme for CD125 of whole bone marrow (A), CD125$^+$ cells backgated into light scatters SSC/FSC (B); LY, lymphocytes; MO, monocytes; GC, granulocytes. CD125 labels 11.9% viable naked mole-rat BM cells, and while normal naked mole rat whole marrow consists of ~24% cells with lymphocytic scatter phenotype (smallest viable nucleated cell fraction), CD125$^+$ cells are enriched to ~92% in the LY scatter population, suggesting a label for B cells as in human blood cells.

Successive gating scheme of viable singlet marrow cells with major cell types gated (C), viable singlet marrow cells histogram for CD34 fluorescence (D), CD34$^+$ cells backgated into SSC/FSC (E). CD34$^+$ cells are predominantly sized between lymphocytic and monocytic leukocyte types, as seen for human HSPCs (Baum et al., 1992).

(F) Postsort with 92 viable events of the sorting strategy for blood CITE-Seq, gate colors referring to Figure 1B.

(G) Filtered blood scRNA-Seq dataset after removing cells with >99% of 0 UMIs (unique molecular identifier; left) and discarding genes expressed in <10 cells (right).

(H) Feature Scatter of filtered and normalized blood scRNA-Seq dataset showing the number of genes per cell vs the number of total counts (UMIs) per cell; cell color referring to Figure 1F.

(I) FRAMA transcriptome mapped to ENSEMBL genome and transcriptome using cds (left) or full transcript sequence (right). Total, total annotated transcripts; mapped, blastn FRAMA exon contig alignment mapped to genomic coordinates (>95% identity, e-value<1e$^{-5}$); mapped 80%, mapping FRAMA exon contig with >80% mapping coverage (percent of contig length covered by blastn alignments).

(J) Structural differences in FRAMA transcripts with a difference in mapping coverage between genome and ENSEMBL transcriptome of >20%; lightblue, transcripts with new exons; blue, transcripts with exon extensions.

(K) Heatmap of the top 10 upregulated genes for each blood cluster, see also Table S4.

(L) Expression of selected genes across blood single-cell clusters; Expression level, scaled UMI counts per cell. Notably, nuclear cell cycle-associated genes (HIRA, NVL), splicing factors (HNRNPD, SRSF7), enzymes (HOGA1, STK36) and hematopoietic phosphatases (DUSP2, CD45) were transcribed at higher levels or in more cells in the TC cluster as compared to CD4- or CD8-TCs.

(M) Bloodcounter leukocyte frequencies (% of total leukocytes); Mean ± s.d. for LY, lymphocytes 47.9±11.7; GR, granulocytes 48.3±12.3; MO, monocytes 3.8±1.8; EO, eosinophils 4.1±4.9; N=31. FACS data Figure 1I: Mean ± s.d. for TC 35.8±8.7; 16±8.9; GC 38.8±13; MO 4.5±1.8; EO 1.9±1.8.

**Figure S2. Mouse Bloodcounter data, Analysis of human and mouse HSPC transcriptomes, Erythroid progenitor quantitation of BM and spleen, Related to Figure 1 and 2.**

Bloodcounter volumetric WBC (A) and platelet (B) numbers across animal age for C57BL/6 mice. $R^2$, Pearson correlation coefficient; p-values were determined by conventional linear regression fitting both slope and intercept; N=66.

(C) Wright staining of naked mole-rat PB film, arrows indicate scored platelets; Scale bar 50µm.

(D) Scoring of PB films. p-values were determined by unpaired Student's t-test; N=10.

Bloodcounter volumetric RBC numbers across animal age for C57BL/6 mice (F). $R^2$, Pearson correlation coefficient; p-values were determined by conventional linear regression fitting both slope and intercept; N[hgl]=31; N[mmu]=66.

(G) Scoring of BM Giemsa sections. p-values were determined by unpaired Student's t-test; N=4.

(H) Viable BM cells stained with FITC-CD90 and PECy7-Thy1.1 give rise to two double positive populations. Backgating as in Figure S1C shows Scatter(GC) gate with 95.2% of cells in the granulocyte scatter properties, Scatter(LY) gate with 88.3% of cells in lymphocytic scatter gate.

(I) Filtered, normalized and log2-transformed CPMs [counts per million] of naked mole-rat BM transcriptomes.

(J) Numbers of differentially expressed genes (DEGs) as determined by limma, see Methods for contrast matrix; log-fold-change cutoff 1; p-value cutoff 0.05.

(K) Human HSPC hierarchy of BLUEPRINT EGAD00001000745 datasets; cell type color used for Figure S1J-K, S6L. HSC, hematopoietic stem cell; CMP, common myeloid progenitor; GMP, granulocyte monocyte progenitor; MEP, megakaryocyte erythrocyte progenitor; MP, megakaryocyte precursor; EB, erythroblast.

(L) Filtered, normalized and log2-transformed CPMs of human CB transcriptomes.

(M) Unsupervised t-SNE clustering. While the unipotent MPs and EBs are clearly separated from stem and progenitors, MEPs and CMPs overlap, resulting in reduced differentially expressed genes at stringent cutoff.

(N) Numbers of differentially expressed genes (DEGs) as determined by limma, see Methods for contrast matrix; log-fold-change cutoff 1; p-value cutoff 0.05.

(O) ssGSEA of human CB HSPCs displaying the top 5 adjusted p-value [q-value] terms from a geneset collection of human and mouse HSPCs (Schwarzer et al., 2017). NES, normalized enrichment score; GeneRatio, (signature ∩ term) / (signature ∩ all terms). Human HSCs match with 5 human HSC terms, human CMPs match with terms MEP, CMP and CD34+, human GMPs match with genesets GMP, human myeloid and murine lymphomyeloid, human MEPs match with terms MEP, CMP and negatively associate with HSC genesets, human MPs match with 3 MK sets and EBs match with 5 erythroid terms. This validates our approach of functional annotation through the geneset collection.

(P) Mouse HSPC hierarchy of GEO GSE74691 datasets; cell type color used for Figure S1O-P, S6L. LT-HSC, long term hematopoietic stem cell; ST-HSC, short term hematopoietic stem cell; CMP, common myeloid progenitor; GMP, granulocyte monocyte progenitor; MEP, megakaryocyte erythrocyte progenitor.

(Q) Filtered, normalized and log2-transformed CPMs of mouse BM transcriptomes.

(R) Unsupervised t-SNE clustering. Note that ST-HSCs cluster within the group range of LT-HSCs, hence will not produce significant differential expression when used with our stringent contrast matrix design, thus we grouped them together as HSCs. Also evident from t-SNE clustering is the central position of CMPs indicating a fluid transition state between HSCs and further committed MEPs and GMPs.

(S) Numbers of differentially expressed genes (DEGs) as determined by limma, see Methods for contrast matrix; log-fold-change cutoff 1; p-value cutoff 0.05. With the applied cutoffs no specific genes for CMPs were detected.

(T) ssGSEA of mouse BM HSPCs displaying the top 5 q-value genesets; each $q < 10^{-7}$. Mouse HSCs match to 4 human HSC and one MK geneset, mouse GMPs are negatively associated with erythroid and lymphoid terms, mouse MEPs are negatively associated with HSC and MLP terms.

If top 5 GeneRatio terms would be displayed, mouse HSCs will match with 2 human and 3 murine HSC terms, mouse GMPs will match with 4 myeloid terms and mouse MEPs will match with human MEP and MK terms, each $q < 10^{-4}$ (data not shown). This exemplifies valid alternative ranking based on GeneRatio or NES to correctly annotate cell types.

Sorting strategy for spleen cells with lineage (LIN) depletion (U) and gating of $LIN^{+}$ CP1 (V); Note the prominent $LIN^{dim}$ population as an upper limit for the $LIN^{-}$ boundary, from which a vertical $LIN^{+}/SSC^{low}$ population sprouts out similar to BM, Figure 2A, featuring lymphocytic cells (data not shown). We thus set the splenic $LIN^{-}$ population to ~35%. Of note, when we exclude $LIN^{dim}$ from $LIN^{-}$ in both BM and spleen to start at ~10% $LIN^{-}$ cells for subsequent gating, CP2 becomes strongly declined while CP1 frequencies did not differ between BM and Spleen (data not shown). In mice the frequency of repopulating, self-renewing stem cells is ~10-fold lower in spleen compared to BM (Morita et al., 2011).

(W) Relative counts of Benzidine-stained cytospins from whole spleen or WBM. p-value determined by Sidak's Two-way ANOVA comparing BM vs spleen between mouse [mmu, N=5] and naked mole-rat [hgl, N=8].

**Figure S3. Cell cycle stainings, Colony assays, Xenograft quantitation, Related to Figure 3.**

(A) Heatmap showing all upregulated genes (top panel) for each CP; Expression, centered and row-scaled voom-transformed CPMs. Numbers of DEGs (bottom panel) as determined by limma, see Methods for contrast matrix; log-fold-change cutoff 1; p-value cutoff 0.05.

(B) Ki-67 / DAPI staining of naked mole-rat WBM, cell cycle stage gates were used for CP1/2.

(C) Pyronin Y / DyeCycle Violet staining of naked mole-rat WBM, cell cycle stage gates were used for CP1/2.

(D) Alkaline Phosphatase (AP) staining of sorted PB cells; scale bar 50µm.

(E) Counts of AP+ cells relative to toal number of cells; N=5.

(F) Benzidine (upper row, BMH), May-Grünwald-Giemsa (middle row, MGG) and AP stainings of picked colonies, see Figure 3D for morphology types; scale bar 50µm. Erythroid CFU and BFU cells contained hemoglobin, the latter at lower frequency than the more mature CFU-E. CFU-M exclusively consisted of large cells with colorless vesicles and a high cytoplasmic/nuclear ratio, while CFU-GM (granulocyte/monocyte) also comprised $AP^+$ cells.

(G) Benzidine-stained colony assays, N=3. Error bars denote s.d., p-value determined by Sidak's Two-way ANOVA.

(H) Untransplanted (irradiated) NSGS host BM stained with naked mole-rat markers; hgl, naked mole-rat cells; mmu, mouse cells.

(I) Transplanted NSGS host BM stained with guinea-pig CD45 and CD45.1; CP2 xenograft at week 4; DN, double negative. The DN fraction contains most $CD34^+$ and $Thy1.1^+/CD34^+$ naked mole-rat cells and a $Thy1.1^-/CD34^-$ population potentially containing xenogenic CP7 (J), the $CD45^+/CD45.1^-$ fraction contains most naked mole-rat GCs [$Thy1.1^{+++}/CD34^-$ in BM] (K), the $CD45.1^+$ fraction features an abundant $Thy1.1^+/CD34^+$ population resembling CP6 in naked mole-rat BM and spleen (L).

(M) Gating strategy to quantify the Thy1.1$^+$/CD34$^+$ double positive cell proportion of each xenograft; hgl, naked mole-rat cells; mmu, mouse cells.

(N) Thy1.1$^+$/CD34$^+$ proportion of engraftment; p-value determined by Tukey's Two-way ANOVA.

**Figure S4. Engraftment subsetting, BM scRNA-Seq, FACS for thymus and lymph nodes, Thymus scRNA-Seq, Related to Figure 3 and 4.**

(A) Gating strategy of NSGS recipient BM using a human CD11b MoAb to selectively label naked mole-rat hCD11b+ cells. CP1 xenografts show ~8-fold less LTCs in the hCD11b– population (B, left) than CP2 xenografts (C, left), whereas hCD11b+/Thy1.1+/CD34+ double positive cell fractions are not different (B, C, right panels).

(D) Postsort QC with 242 viable events of the sorting strategy for BM CITE-Seq, gating refers to Figure 2D.

(E) Signal intensities of CD11b of gates from S2D; CD34++/Thy1.1++, CP1/2 [steelblue AUC=area under curve].

(F) Expression of BC cluster marker genes associated with BC functions in human and mouse, see also Table_SX_BM_markers.

(G) Blendplot showing BM TARP vs CYTL1.

(H) Blendplot showing BM GP9 vs CLEC1B; Note the exclusive overlap at the HSPC4 cluster region without GATA2 expression, see also Figure 4D.

(I) Blendplot showing BM VPREB1 vs VPREP3; Note the BC2 cluster as exclusive overlap of VPREB1 and VPREB3.

(J) Micrographs from the cervical region of a naked mole-rat, showing the superficial (left) situs of thyroid (blue arrowhead) and parathyroid glands (black arrowheads); The profound (right) situs with several lymph nodes (white arrowheads).

(K) Hematoxylin & Eosin staining of a parathyroid from the superficial cervical region of a naked mole-rat, scale bar 100µm.

(L) Frequencies of mature (left plot) and progenitor (right plot) populations across lymphatic organs. THY, thymus, N=31; B lymph node, N=16; T lymph node, N=7.

FACS gating of CD125+ BCs in BM (M), spleen (N) and B lymph node (O). Note that the B lymph node CD125+ population is split into two separate CD125+ and CD125++ fractions, possibly due to an activated state. This was seen in 1/5 animals we co-stained BLNs for CD125, with no apparent disease condition noticed elsewhise, other animal BLN showed CD125+ staining pattern comparable to BM.

(P) Frequencies of CD125+ in BM (N=8), Spleen (N=7) and B lymph nodes (LN, N=10); p-value determined by Dunnett’s One-way ANOVA.

(Q) Postsort with 102 viable events of the sorting strategy for thymus CITE-Seq, gating refers to Figure 4G. Although thymic LTCs comprised 14% of total thymocytes sorted, LTC markers (HOXA9, ICAM2, CYTL1, GATA2, ID2, C5, AMN1) were not differentially expressed (TableS2, S4), arguing for a thymus-seeding LTC subpopulation different from BM LTCs and co-clustering with TSPs.

(R) CITE-Signals for thymus single cell clusters, data centered and scaled.

(S) Blendplot showing thymus ID3 vs NOTCH1 expression; framed region same as in Figure 5K-L.

**Figure S5. ScRNA-Seq combined dataset, pseudotime, Related to Figure 5.**

(A) Proportions of sorted cell fractions pooled for Cite-Seq libraries as measured from each tissues postsort analysis.

(B) Proportions of Louvain clusters using the combined BM, PB and thymus libraries.

UMAP dimensionality reduction showing the 19 clusters (C) or the tissue library origin (D) of the dataset. The sensitivity of clustering resolution was outlined by emergence of recurring clusters <51 cells, i.e. PCs from Figure 4A reappear as BM.PCs with the same markers (Table S4; see Methods). Similarly, 20 stromal cells (SC, Figure 4A) were reclustered in the joint dataset as 23 BM.SCs overexpressing markers such as Type I Collagens, Osteocalcin, Osteonectin, Bone Sialoprotein 2 or IFITM5 with the highest fold-changes across the entire data (Table S4).

UMAP highlighting EPCAM (E), CEBPα (F), CEBPε (G), IL7R (H) or TARP (I) expression throughout the trajectory; Expression, log2 UMI counts.

(J) UMAP dimensionality reduction as blueprint for MST trajectories, see legend for color code. Connecting line, trajectory; Circles, trajectory milestones; Arrowheads, pseudotime direction; coloring by nearest cell.

(K) Dendrogram of MST pseudotime with Louvain clustering input, root set to a seed HSC cluster cell.

Diffusion map (L) or Ouija (M) pseudotime (x-axis) for indicated genes (left y-axis), cells ordered into Louvain cluster categorials (right y-axis); Expression curves smoothed using GAM models. Note that the density kernels of TSP/ETP/PB.TC did not form a "waterfall" or "stairway" pattern indicating successive developmental stages with Diffusion map pseudotime, as opposed to *Ouija*; The same was seen for BM.MY/PB.DC/PB.MO/PB.GC. *Ouija* handles the myeloid branch BM.MY/PB.DC/PB.MO/PB.GC more consistent than PAGA and MST by splitting BM.My and assigning a smaller cluster proportion with a density kernel analogous to BCP, hence both anchoring at MPP. Interestingly, the stairway from MEP to ERY4 is more accentuated with

Diffusion Map pseudotime. Though the major problem for all trajectory inference algorithms used in our study remained the bifurcation of lymphopoiesis at the BM BCP stage integrating thymic TC fractions with PB TC and BC clusters. Future studies addressing this issue are warranted.

**Figure S6. Ki-69 staining, FACS quantifications, 3-species gene expression, Rhodamine efflux, Related to Figure 6.**

(A) Filtered, normalized, log2-transformed and median-centered TPMs [transcripts per million] of human, mouse and naked mole-rat HSPC transcriptomes, color code see Figure S6E.

(B) Representative gating strategy for mouse $LIN^{–}$/$Sca\text{-}1^{+}$/$Kit^{+}$ (LSKs); LIN gate set to 10% viable cells in a biplot analog to Figure 2A.

(C) Representative gating strategy for human $LIN^{–}$/$CD38^{lo}$/$CD34^{+}$ (HSCs); LIN gate set to 10% viable cells in a biplot analog to Figure 2A.

(D) Cell cycle staining with Ki67 (CP1/LTC, N=12); young mouse BM (N=4) was used for LK and LSK. p-value determined by Tukey's Two-way ANOVA. Note that CP1 cells do not differ in any cell cycle stage from LKs.

(E) Expression levels of HSPC transcriptomes across 3 species, log2 TPMs.

(F) Representative gating of naked mole-rat BM for viable TCs, GCs, ERY ($Thy1.1^{–}$/$CD34^{+/++}$) and $Thy1.1^{++}$/$CD34^{++}$ (T34), no LIN was applied.

(G) MitoTracker signal across cell types in naked mole-rat BM; merged analysis, N=5. Mitochondrial load increased in erythroid cells (ERY) and decreased in TCs.

(H) Mean Fluorescence Intensity (MFI) of MitoTracker measurements in mouse (N=4), human (N=4) and naked mole-rat (N=5) BM. p-value determined by Tukey's One-way ANOVA.

(I) MFI of MitoSOX measurements in mouse (N=4) and naked mole-rat (N=5) BM. p-value determined by Tukey's One-way ANOVA.

(J) MFI of CellROX measurements in mouse (N=4), human (N=4) and naked mole-rat (N=5) BM. p-value determined by Tukey's One-way ANOVA. Concordantly, oLSKs with less active mitochondria as shown by MSO have less ROS levels than yLSKs.

(K) Merged JC-1 fluorescence biplots for unstained (top left), ERY (bottom left), TC (top right) and GC (bottom right) naked mole-rat BM cells gated according to Figure S7D.

(L) Proportions of JC-1 MMP populations gated from Figure 7J-L in mouse (N=4), human (N=4) and naked mole-rat (N=5) BM. p-value determined by Sidak's Two-way ANOVA.

(M) MFI of Aldefluor measurements; N=6. p-value determined by Tukey's One-way ANOVA.

(N) Rhodamine 123 efflux; merged analysis samples from (O) MFI of Rhodamine efflux measurements in mouse (N=4) and naked mole-rat (N=5) BM. p-value determined by unpaired Student's t-test.

**Figure S7. Side Population staining, summary table, Related to Figure 6.**

(A) Side population (SP) staining using DyeCycle Violet (DCV) fluorescence on WBM. LSKs are backgated into SSC/FSC properties. When the $SP^{lo}$ population is crossgated into Sca-1/Kit biplot, > 90% of $SP^{lo}$ cells are LSKs (data not shown).

(B) Mouse $SP^{lo}$ cells are backgated into FSC/SSC, where they appear as a single population of low granularity sized between lymphocytes and monocytes (top panels). Naked mole-rat $SP^{lo}$ cells are backgated into FSC/SSC they appear as a single population of low granularity and sized between lymphocytes and monocytes (bottom panels). Note that the naked mole-rat $SP^{hi}$ is far less abundant than in mice, and although co-staining with Propidium Iodide removed dead cells, the SP "leaks" into an autofluorescent streak, signs for an unspecific SP effect we also detected with rat and guinea pig WBM (data not shown). Data obtained using 90min cell incubation @ 37°C, similar results for middle and bottom panels were obtained with reduced incubation time to 45min (data not shown).

(C) Expression levels of orthologs involved in HIF1α respiration pathway; scale, TPM z-score.

(D) Summary of homeostatic hematopoiesis between 3 species. Mouse PB cell data taken from (STEMCELL Technologies, DOCUMENT #28048 | VERSION 3.0.0). Mouse Mean PLT Figure S2B, $1235 \pm 520e^{9}/l$; Mouse Mean RBC Figure S2F, $9.1 \pm 0.6e^{12}/l$. Human PB cell data taken from (STEMCELL Technologies, DOCUMENT #23629 | VERSION 4.0.0).
