## Supplementary material for "The hematopoietic landscape at single-cell resolution reveals unexpected stem cell features in naked mole-rats": Table S3 scRNA-Seq BM partitions GSEA

**Table S3. fGSEA of 4 partitions derived from 16 clusters of BM scRNA-Seq, Related to Figure 4**

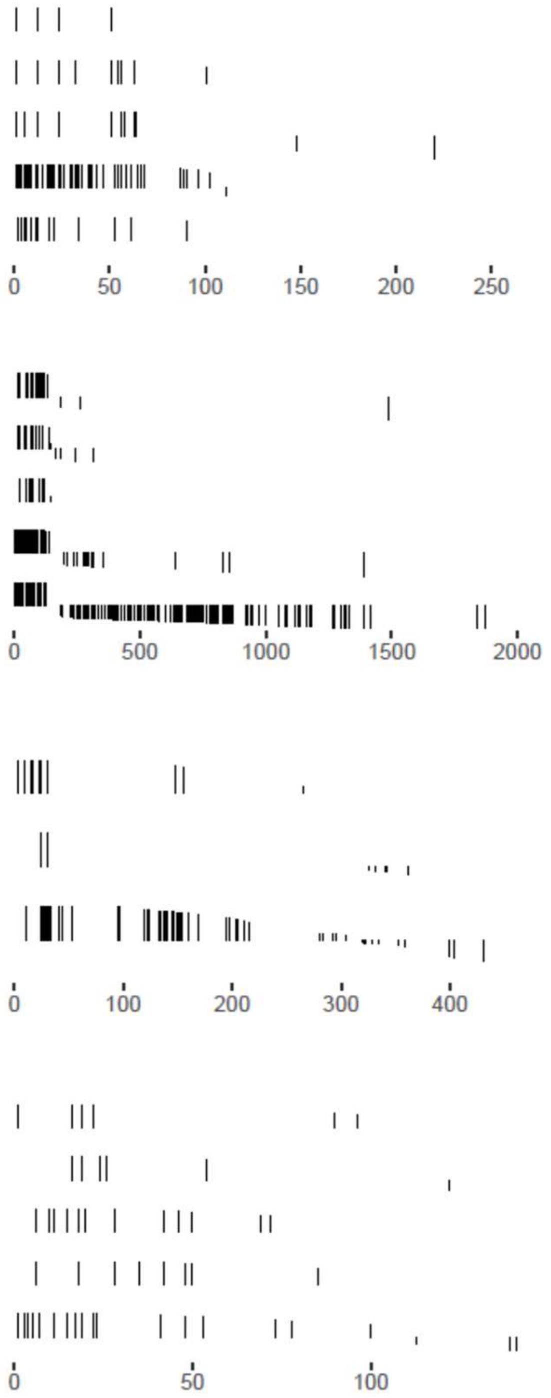


| | pathway | gene ranks | NES | p.adj |
|---|---|---|---|---|
| HSPC | EBERT_hHSC | | 1.87 | 4.7e−02 |
| | EBERT_hCD34+ | | 2.30 | 3.5e−03 |
| | KLUSMANN_NCODE_hCD34+ | | 2.20 | 1.0e−02 |
| | EMMRICH_nLYMPHOMYELOID_UP | | 4.24 | 4.7e−04 |
| | EMMRICH_nLTC_UP | | 2.72 | 4.7e−04 |
| ERY | EBERT_hERYTHROID | | 5.22 | 4.5e−04 |
| | JAATINEN_hHSC_DN | | 4.18 | 4.5e−04 |
| | ROSSI_mERYTHROID | | 4.28 | 4.5e−04 |
| | EMMRICH_nERYTHROID_UP | | 7.24 | 1.1e−03 |
| | EMMRICH_nEEP_UP | | 4.21 | 5.9e−03 |
| BC | ROSSI_mBC | | 1.68 | 2.9e−02 |
| | LAURENTI_hHSC | | 1.75 | 2.5e−02 |
| | EMMRICH_nLYMPHOMYELOID_UP | | 1.47 | 2.9e−02 |
| MY | GEORGOPOLOUS_GMPvsCD33+ | | 1.60 | 1.3e−01 |
| | GEORGOPOLOUS_MYELOID | | 1.46 | 2.7e−01 |
| | KLUSMANN_Arraystar_hMYELOID | | 1.58 | 1.3e−01 |
| | ROSSI_mGRAN | | 1.50 | 2.3e−01 |
| | LAURENTI_hGMP | | 1.78 | 1.9e−02 |
