## Supplementary material for "The hematopoietic landscape at single-cell resolution reveals unexpected stem cell features in naked mole-rats": Table S6 BM PB THY clusters GSEA

**Table S6. fGSEA of 15 clusters of combined BM/PB/THY libraries, Related to Figure 6**

| Cluster | pathway | NES | pval |
|---|---|---|---|
| HSC | EBERT_hHSC | 2.29 | 5.0e−05 |
| HSC | EBERT_hCD34+ | 2.39 | 5.1e−05 |
| HSC | KLUSMANN_NCODE_hCD34+ | 2.09 | 1.3e−03 |
| HSC | EMMRICH_nLYMPHOMYELOID_UP | 3.90 | 2.9e−05 |
| HSC | EMMRICH_nLTC_UP | 2.81 | 2.6e−05 |
| MPP | EMMRICH_nLYMPHOMYELOID_UP | 2.19 | 3.4e−05 |
| PB-DC | EBERT_hMYELOID | 1.56 | 2.4e−03 |
| PB-DC | KLUSMANN_Arraystar_hMONO | 1.64 | 4.1e−04 |
| PB-DC | EMMRICH_nLYMPHOMYELOID_UP | 1.65 | 1.2e−04 |
| PB-MO | EBERT_hMYELOID | 2.50 | 1.2e−05 |
| PB-MO | JAATINEN_hHSC_DN | 1.96 | 2.5e−05 |
| PB-MO | KLUSMANN_Arraystar_hMONO | 2.12 | 1.3e−05 |
| PB-MO | KLUSMANN_Arraystar_hMYELOID | 2.03 | 1.4e−05 |
| PB-MO | KLUSMANN_NCODE_hMONO | 2.03 | 2.7e−05 |
| PB-MO | LAURENTI_hGMP | 1.84 | 4.3e−04 |
| PB-MO | EMMRICH_nLYMPHOMYELOID_UP | 2.30 | 1.2e−05 |
| PB-MO | EMMRICH_nMMPP_UP | 1.88 | 2.2e−04 |
| PB-GC | EBERT_hMYELOID | 4.03 | 1.6e−05 |
| PB-GC | GEORGOPOLOUS_earlyLYMPHOMYELOID | 2.58 | 1.7e−05 |
| PB-GC | JAATINEN_hHSC_DN | 2.68 | 1.7e−05 |
| PB-GC | KLUSMANN_Arraystar_hGRAN | 3.37 | 1.7e−05 |
| PB-GC | KLUSMANN_Arraystar_hMYELOID | 3.64 | 1.6e−05 |
| PB-GC | KLUSMANN_NCODE_hGRAN | 3.19 | 1.7e−05 |
| PB-GC | ROSSI_mGRAN | 3.46 | 1.7e−05 |
| PB-GC | EMMRICH_nLYMPHOMYELOID_UP | 3.72 | 1.5e−05 |

gene ranks

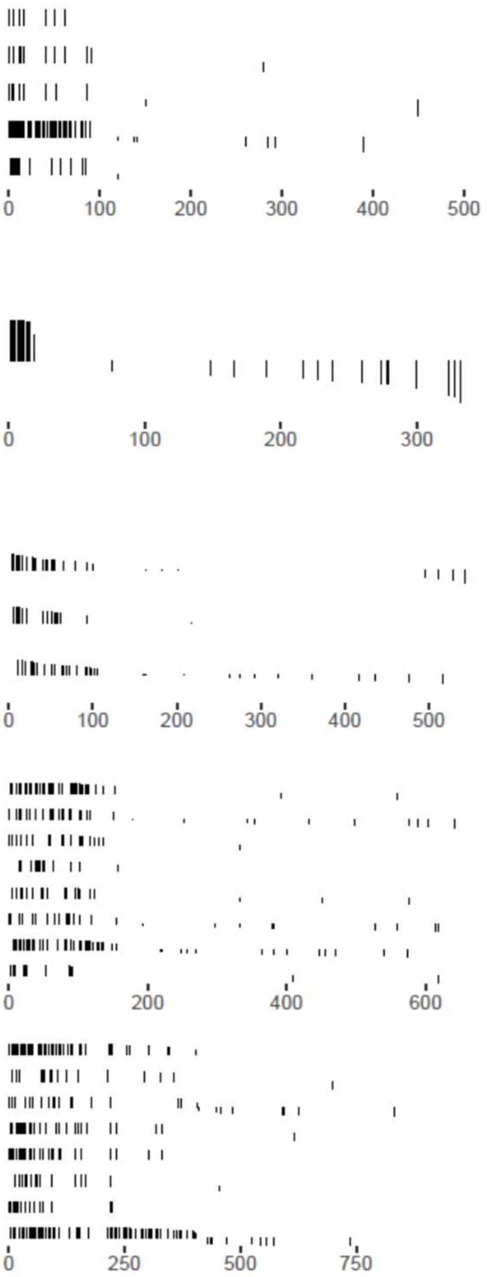

**Table S6, continued**

| | pathway | gene ranks | NES | pval |
|---|---|---|---|---|
| BCP | GEORGOPOLOUS_proBC | | 1.66 | 9.9e−04 |
| | KLUSMANN_Arraystar_hBC | | 1.59 | 2.8e−03 |
| | KLUSMANN_Arraystar_HSC_CB vs ALL_Top200 | | 1.59 | 3.6e−03 |
| | KLUSMANN_NCODE_hBC | | 1.54 | 1.4e−03 |
| | LAURENTI_earlyB | | 1.68 | 1.2e−03 |
| TSP | EBERT_hLYMPHOID | | 2.24 | 2.1e−05 |
| | EBERT_hTC | | 2.24 | 2.1e−05 |
| | KLUSMANN_Arraystar_hTC | | 1.84 | 1.9e−03 |
| | ROSSI_mTC | | 2.24 | 2.1e−05 |
| | LAURENTI_hETP | | 2.39 | 2.0e−05 |
| | REGEV_G1−S_Core Set | | 1.79 | 1.2e−03 |
| ETP | ROSSI_mTC | | 2.31 | 2.8e−05 |
| | LAURENTI_hETP | | 2.33 | 2.9e−05 |
| PB-TC | EBERT_hLYMPHOID | | 2.49 | 3.5e−05 |
| | EBERT_hTC | | 2.49 | 3.5e−05 |
| | KLUSMANN_Arraystar_hTC | | 2.46 | 3.6e−05 |
| | ROSSI_mTC | | 2.65 | 1.7e−05 |
| | GSE15330_mHSCvsMEP_UP | | 2.03 | 9.6e−04 |
| BC | EBERT_hBC | | 2.98 | 2.3e−05 |
| | GEORGOPOLOUS_proBC | | 1.99 | 3.9e−03 |
| | GEORGOPOLOUS_lateLYMPHOMYELOID | | 1.96 | 1.3e−04 |
| | GOODELL_mBC | | 2.15 | 9.7e−04 |
| | KLUSMANN_Arraystar_hBC | | 3.57 | 2.3e−05 |
| | KLUSMANN_NCODE_hBC | | 2.64 | 2.2e−05 |
| | ROSSI_mBC | | 3.37 | 2.3e−05 |
| | EMMRICH_nLYMPHOMYELOID_UP | | 3.06 | 2.3e−05 |

**Table S6, continued**

| | pathway | gene ranks | NES | pval |
|---|---|---|---|---|
| ERY1 | EMMRICH_nERYTHROID_UP | | 3.12 | 3.9e−05 |
| | EMMRICH_nEEP_UP | | 3.15 | 8.3e−05 |
| ERY2 | KLUSMANN_Arraystar_hERYTHROID | | 1.95 | 1.1e−03 |
| | LAURENTI_hETP | | 2.35 | 1.6e−05 |
| | REGEV_G1−S_Core Set | | 2.16 | 8.9e−05 |
| | REGEV_G2−M_Core Set | | 2.76 | 1.6e−05 |
| | EMMRICH_nERYTHROID_UP | | 2.84 | 1.4e−05 |
| | EMMRICH_nEEP_UP | | 2.69 | 1.4e−05 |
| ERY3 | EBERT_hERYTHROID | | 3.70 | 1.6e−05 |
| | KLUSMANN_NCODE_hERYTHROID | | 2.35 | 2.5e−04 |
| | ROSSI_mERYTHROID | | 3.60 | 1.7e−05 |
| | LAURENTI_hMEP | | 2.19 | 1.3e−03 |
| | GSE35685_hHSCvsCD34+CD10+_UP | | 2.04 | 3.0e−03 |
| | EMMRICH_nERYTHROID_UP | | 4.24 | 1.5e−05 |
| | EMMRICH_nEEP_UP | | 1.87 | 6.4e−03 |
| ERY4 | EBERT_hERYTHROID | | 1.86 | 1.6e−04 |
| | EMMRICH_nEEP_UP | | 1.52 | 2.2e−04 |
| BM.SC | EMMRICH_nEEP_UP | | 1.28 | 3.0e−04 |

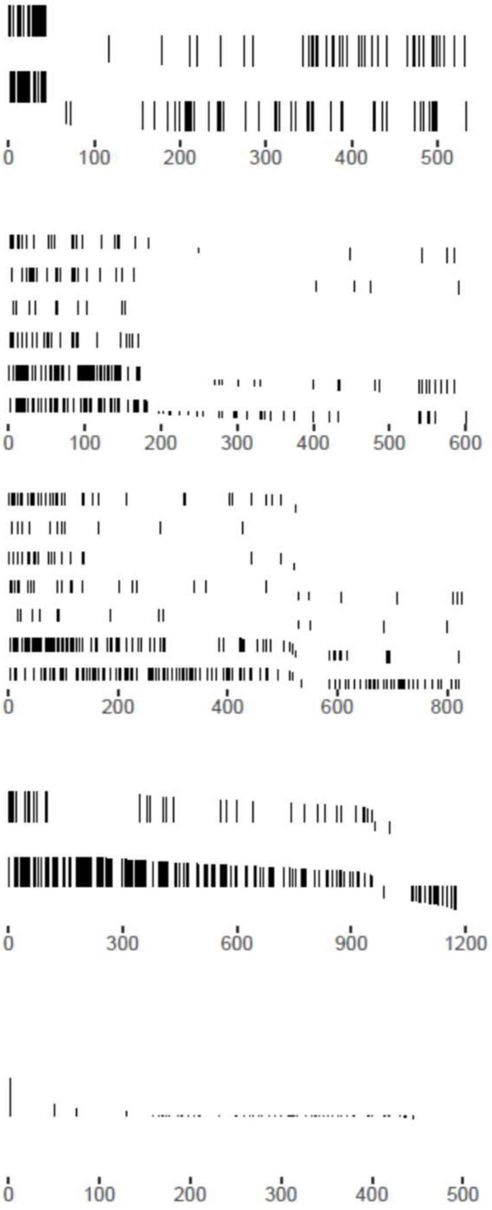
