## Supplementary material for "The hematopoietic landscape at single-cell resolution reveals unexpected stem cell features in naked mole-rats": Table S7 KEGG pathway OXPHOS

### Table S7. KEGG pathway OXPHOS highlighting differentially expressed genes in naked mole-rat HSPCs, Related to Figure 7

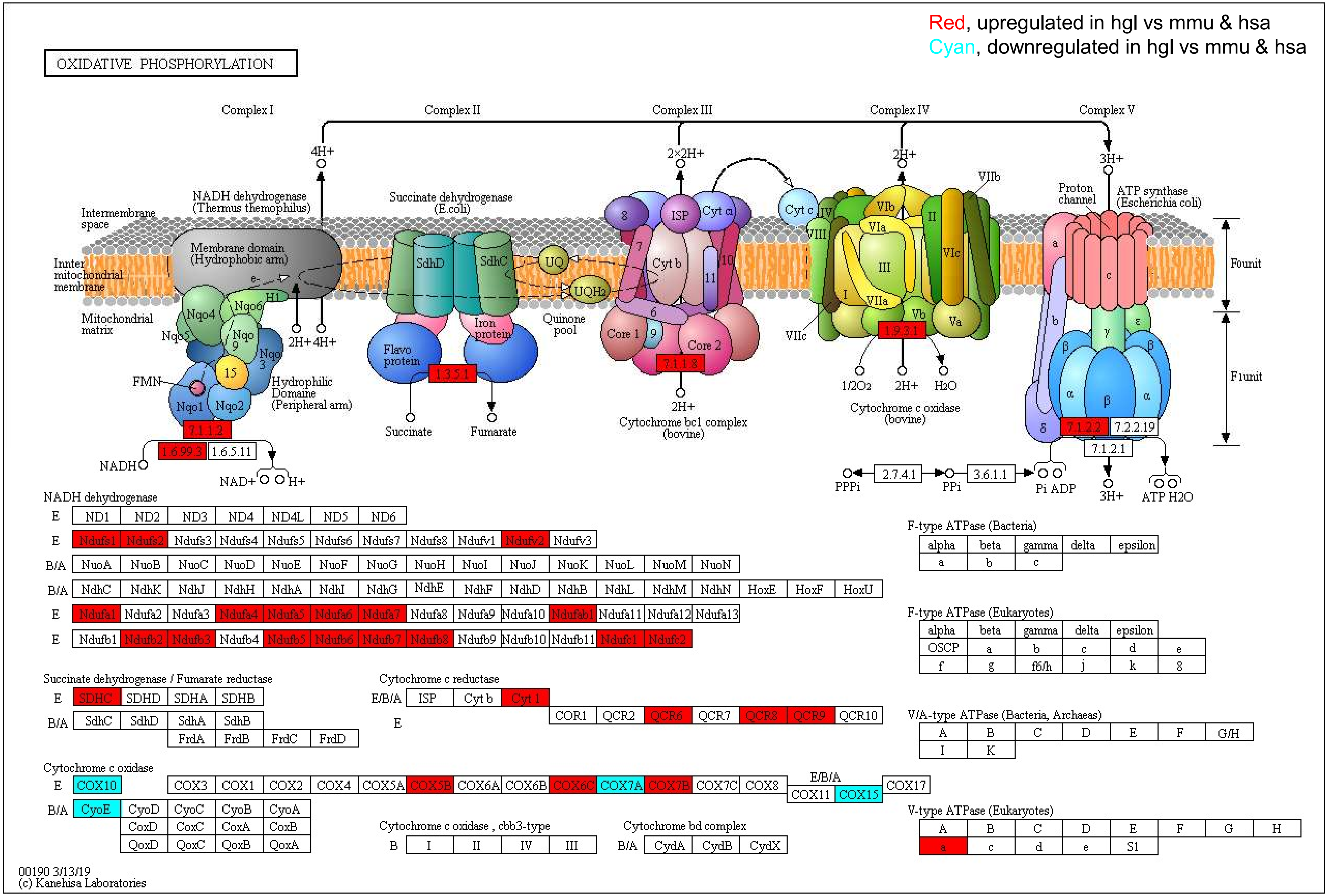
